# Targeting RhoA activity rejuvenates aged hematopoietic stem cells

**DOI:** 10.1101/2025.04.23.647902

**Authors:** Eva Mejía-Ramírez, Pablo Iañez Picazo, Barbara Walter, Sara Montserrat-Vazquez, Francesco Affuso, Stefan Wieser, Fabio Pezzano, Loïc Reymond, Jorge Castillo-Robles, Francesca Matteini, Loris Mularoni, Dídac Maciá, Ángel Raya, Verena Ruprecht, Yi Zheng, Paula Petrone, M. Carolina Florian

## Abstract

Biomechanical alterations contribute to the decreased regenerative capacity of hematopoietic stem cells (HSCs) upon aging. RhoA is a key regulator of mechano-signaling but its role for mechanotransduction in stem cell aging has not been investigated yet.

Here, we show that murine HSCs respond to increased nuclear envelope (NE) tension by inducing NE translocation of P-cPLA2, which cell intrinsically activates RhoA. Interestingly, aged HSCs experience physiologically higher intrinsic NE tension, associated with increased NE P-cPLA2 and RhoA activity. Reducing RhoA activity lowers NE tension in aged HSCs. Feature image analysis of HSC nuclei reveals that chromatin remodeling is associated to RhoA inhibition, which includes the restoration of youthful levels of the heterochromatin marker H3K9me2 and a decrease in chromatin accessibility and transcription at retrotransposons. Eventually, we demonstrate that RhoA inhibition upregulates Klf4 expression and transcriptional activity, improving aged HSCs regenerative capacity and lympho/myeloid skewing *in vivo*. Overall, our data support that an intrinsic mechano-signaling axis dependent on RhoA can be pharmacologically targeted to rejuvenate stem cell function upon aging.

## INTRODUCTION

Aging is characterized by the decline in tissue function and is the primary risk factor for major diseases. In particular, aged HSC functional decline critically impacts not only on their ability to regenerate the hematopoietic system and to support lymphoid cell production over time^1–4^ but also directly contributes to major aging-related diseases^5,6^. Aging is a complex, multifaceted process that is accompanied by biomechanical changes affecting tissues and organs and also cells and subcellular organelles^7–12^. Among others, these biomechanical changes include alterations in NE tension and aging correlates with progressive changes in the nucleus mechanical integrity and impaired mechanotransduction^11,12^. However, how to possibly target changes in nuclear mechano-signaling to explain and prevent aging of somatic stem cells is still largely under-investigated.

In addition, epigenetic alterations are considered one of the primary hallmarks of aging^13^ and despite the large amount of data demonstrating the occurrence of an epigenetic drift upon stem cell aging and disease, there is a lack of knowledge on molecular mechanisms to explain this epigenetic drift and whether it possibly associates to mechanical alterations of the nuclear and chromatin architecture.

Mechanical forces trigger multiple signaling pathways that converge in the activation of RhoA^14^, which is a small RhoGTPase that can cycle between an active (RhoA-GTP) and an inactive (RhoA-GDP) status. RhoA is a key regulator of mechanotransduction^15^ regardless of whether the activating mechanical stimulus is cell extrinsic, as occurs in cells responding to alterations of substrate stiffness^16^, or cell intrinsic, like for example when the cell nucleus acts as a mechanosensor of genomic changes^17–19^. So far, in HSCs RhoA has been shown to be important for cytokinesis^20^. *In vivo,* knocking out RhoA in bone marrow cells induces a dramatic phenotype, characterized by a multilineage hematopoietic failure due to programmed necrosis of hematopoietic progenitors, while HSCs retain long-term engraftment potential but fail to produce hematopoietic progenitors and lineage-defined blood cells^20^.

Here, we investigate the role of RhoA in nuclear mechanotransduction in HSCs and its involvement in preserving nuclear architecture and stem cell function upon aging. Our data reveal that RhoA activity increases upon increased NE tension in HSCs, which is intrinsically altered upon aging and can be targeted to improve *in vivo* function of aged blood stem cells.

## RESULTS

### RhoA is necessary for HSCs to survive under increased NE tension

The nuclear membrane is able to stretch under various pathophysiological conditions involving nuclear lamina weakening, which include aging and laminopathies^21^, and stretching of the nucleus is thought to be a fundamental mechanism engaging nuclear mechnotransduction^21,22^. Therefore, we wondered if nuclear stretching might be engaging RhoA mechano-signaling pathways in HSCs. HSCs are notoriously small sized non adherent cells with an average cell diameter of 10 µm and a high nuclear/cytosol ratio. To induce nuclear stretching in HSCs, we took advantage of a previously established cell confinement device^18,23^ (**Figure S1A**). We isolated HSCs (sorted as Cd11b^-^, Ter119^-^, Cd8^-^, Cd5^-^, B220^-^, Gr1^-^, c-kit^+^, Sca1^+^, Flt3^-^, CD34^-^) from the bone marrow (BM) of young adult (8 to 20 weeks old) C57Bl6 mice and we seeded them on fibronectin-coated coverslips followed by confinement with non-adhesive coverslips functionalized with pillars of 8, 5 and 3 µm height for 2 hours (**Figure 1A** and **Figure S1A-B**). Upon confinement, HSCs were fixed and stained with DAPI (4′,6-diamidino-2-phenylindole) to image and measure the nuclear stretching, and with an anti-RhoA-GTP antibody for quantifying the activation levels of RhoA. By 3D single-cell confocal microscopy, we observed that cell confinement produced a reduction of the nuclear height proportional to the applied level of confinement (**Figure 1B**). The maximum nuclear diameter increased significantly already under 8 µm confinement, and it reached its maximum at almost 10 µm when HSCs were confined at 5 µm, showing a significant inverse correlation with nuclear height (**Figure 1B**). We observed the formation of nucleoplasm containing blebs, which was progressive with confinement and particularly prominent when HSCs were confined at 5µm (**Figure 1C**). This can be quantified looking at the Excess of Perimeter (EOP) of the largest 2D slide^24^ and at the major axis length, which significantly increased in HSCs under the 5 µm confinement conditions (**Figure S1C-D**). Interestingly, nuclear stretching was paralleled by a progressive alteration of the DAPI intensity profile of the nucleus and by a sharp increase in the activation levels of RhoA, which was mildly but directly correlated with the increase in nuclear diameter (**Figure 1C** and **Figure S1E**). Indeed, RhoA-GTP levels increased significantly in direct correlation to the level of confinement at 8 µm and 5 µm (**Figure 1C**). When the nuclear height was reduced to 3 µm, RhoA-GTP levels decreased dramatically. In this condition of tight confinement, HSCs displayed severe nuclear rupture, which suggests cell distress and the inability to react against such a reduction of nuclear height, as evidenced by the dramatic reduction of the nuclear volume and DAPI intensity profile, the appearance of nuclear blebs and the extreme standard deviation variability of the EOP (**Figure 1C** and **Figure S1C-E** and **Video S1-4**). This data is consistent with a threshold of up to 70% compression of the nuclear height for HSC survival and consequent RhoA activation before irreversible nuclear lamina rupture, in agreement with what was described for other cells^24^.

**Figure 1.**
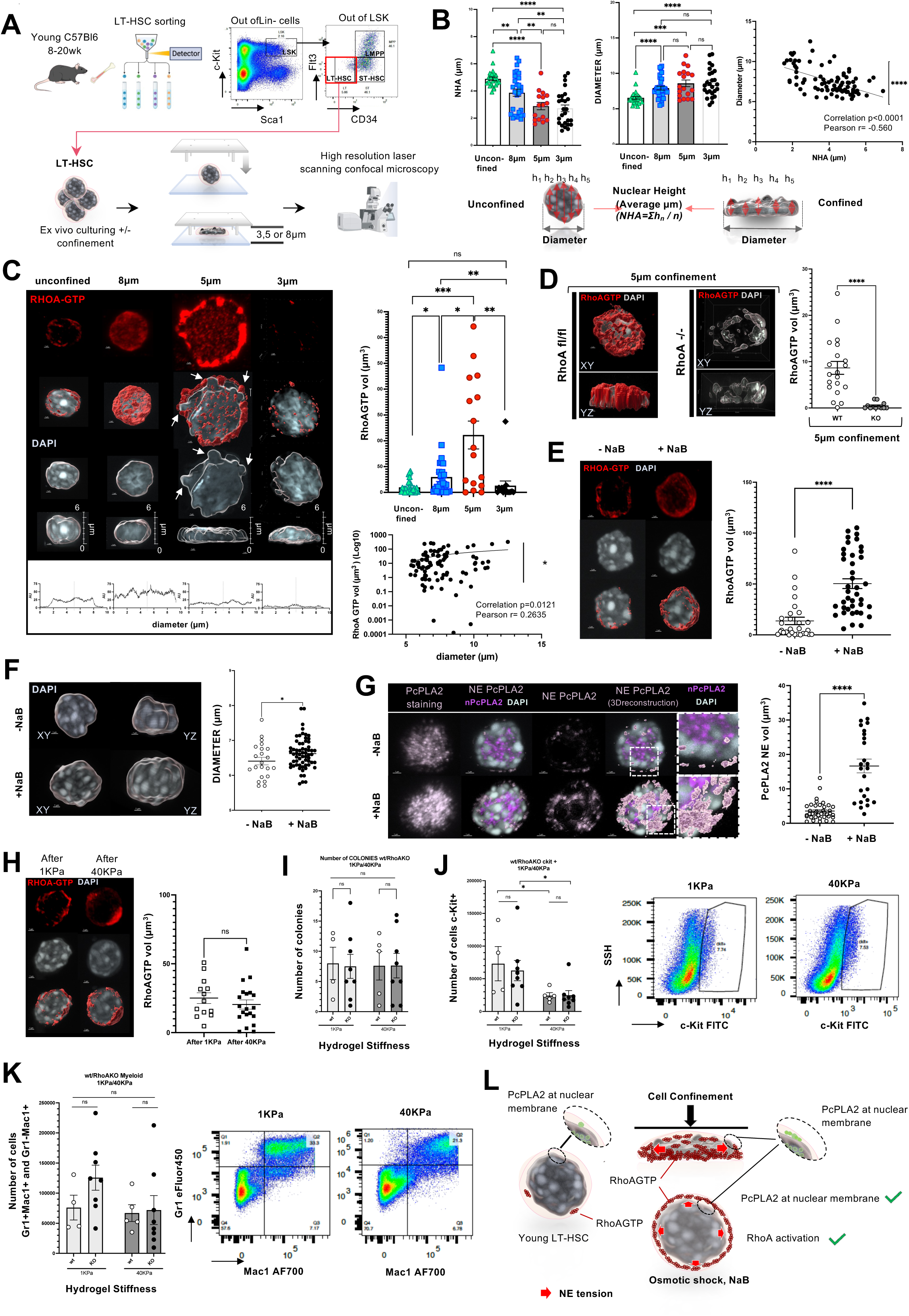
Increasing nuclear envelope (NE) tension activates RhoA in hematopoietic stem cells (HSC). **A,** Representative HSC gating strategy and experimental set-up for the confinement experiments. **B.** Two representative nuclei are depicted to illustrate the strategy to measure the Nuclear Height Average (NHA) and diameter. NHA is calculated as the sum of *n* measurements of the nucleus height in the YZ axis, divided by the *n* number of measurements. Nuclear Height Average (NHA) and Diameter are represented against the different confinement conditions used in the experiments and a correlation plot between both is shown from 3 independent experiments. Data on graphic bars have been analyzed using Mann-Whitney two-tailed tests **p<0.01; ***p<0.001, ****p<0.0001. Data on correlation plot have been analyzed by simple linear regression. Pearson r coefficient and P value are shown. ****P<0.0001 **C.** Representative images of 3D immunofluorescence reconstruction from XY and YZ axes of single HSCs under different levels of confinement of 3 independent experiments. Antibody anti-RhoAGTP was used to stain active RhoA (red). The nuclei are stained by DAPI (gray). Scale bars=1 µm. Movies of these confocal acquisitions are available in **Video S1-4**. Quantification of RhoAGTP (volume of positive signal by intensity) under different confinements: unconfined, 8 µm, 5 µm and 3 µm. Mann-Whitney-test, two-tailed *n*=3\**p<0.05; **p<0.01, ***p<0.001*. Correlation plot in between the RhoAGTP and the diameter is shown. Data on correlation have been analyzed by simple linear regression. Data is obtained from 3 independent experiments. Pearson r coefficient and P value are shown. \**p<0.05*. **D,** Representative images of 3D confocal reconstruction showing LT-HSC from RhoA^fl/fl^ and RhoA^-/-^ mice. Cells were treated *in vitro* with 4OH-tamoxifen o/n and then confined under 5 µm for 2 hours. Cells were stained for RhoAGTP (red) and DAPI (gray). Graph shows volume in µm^3^ of RhoAGTP signal. n=3 Mann-Whitney two-tailed, ****p<0.0001 **E,** Representative images of 3D confocal reconstruction showing RhoAGTP signal (red) and DAPI (gray) of HSCs treated with NaB. Graph shows volume in µm^3^ of RhoAGTP signal. n=3, Mann-Whitney two-tailed \*\*\*\**p*<0.0001. **F,** Representative images of 3D confocal reconstruction showing DAPI staining of nuclei from young control LT-HSC and young treated with NaB. Graph shows the maximum diameter at XY in µm at each condition. n=3, Mann-whitney test one-tailed \**p*<0,05. **G,** Representative images of 3D confocal reconstruction showing phospho-cPLA2 (PcPLA2) (pink) and DAPI (gray) in young control and young NaB treated LT-HSCs. By using image analysis software (Imaris and Volocity) we have compartmentalize PcPLA2 signal within the nucleus and at the nuclear envelope (NE). We have assigned light pink to NE PcPLA2 and magenta to nuclear PcPLA2. NE PcPLA2 only is also shown. The zoom inset shows membrane localization of PcPLA2 which has been reconstructed in light pink in the last panel. Graph showing the volume in µm^3^ of PcPLA2 signal at the NE. Movies of representative confocal acquisitions are available in **Video S5-6**. n=3, Mann-Whitney, two-tailed \*\*\*\**p*<0.0001. **H,** Representative images of 3D confocal reconstruction showing HSC after 16h incubation on hydrogels of 1 KPa or 40 KPa stiffness, stained with RhoAGTP (red) and DAPI (gray). Graph shows volume in µm^3^ of RhoAGTP signal. n=2; Mann-Whitney, two-tailed. **I,** Graphs showing the number of colonies and the number of cells on CFU assays of wt and RhoAKO HSCs after incubation on hydrogels of different stiffness. n=4, Mann-Whitney,two-tailed. Scale bars=1µm. **J and K,** Graphs showing haematopoietic progenitor cells (c-Kit+) (J) and myeloid cells (mac1+gr1+ and mac1+) (K) in CFU assays of Rhoa^fl/fl^ and RhoA^-/-^ LT-HSC after incubation on hydrogels at 1KPa or 40KPa. n=4, Mann-Whitney, two-tailed, \**p*<0.05. **L,** Graphics depicting RhoAGTP and PcPLA2 under normal conditions and nuclear stretching, either by confinement or NaB treatment.

To investigate whether RhoA is necessary for the response to nuclear deformation in HSCs, we isolated HSCs from *CreERt2*X*RhoA^flox/flox^* mice, in which the activity of the Cre recombinase to knock-out *RhoA* can be induced *in vitro* by overnight treatment with 4-hydroxy tamoxifen (**Figure S1F;** hereafter referred to as RhoA knock-out (KO) HSCs). To note, after 12-16 hours from induction of the Cre-recombinase, levels of RhoA-GTP were clearly reduced (**Figure S1G**). Next, we confined the cells at 5 µm, according to the protocol described above to image nuclei and quantify the impact of RhoA knock-out. Surprisingly, in RhoAKO HSCs, the nuclei were completely broken, indicating that RhoAKO HSCs were unable to resist the 5 µm confinement (**Figure 1D**).

Since our experiments were performed *in vitro* on sorted non adherent HSCs, we reasoned that RhoA activation might be intrinsically induced by the mechanical tension at the nuclear envelope (NE). To investigate this hypothesis, we decided to quantify the phosphorylated form of the nuclear protein cPLA2 (P-cPLA2), which translocates to the NE due to a physical process mediated by tension at the NE^21,22,25,26^. NE P-cPLA2 catalyzes the hydrolysis of phospholipids releasing arachidonic acid (AA)^18,22,23,27,28^, which is a well-established activator of RhoA^29,30^. We stained P-cPLA2 and quantified its NE localization, which increased when HSCs are under confinement, consistent with the increased RhoA activation (**Figure S1H**).

To further investigate whether intrinsic RhoA activation is induced by NE tension, we treated freshly sorted HSCs with sodium butyrate (NaB), a histone deacetylase inhibitor known to increase levels of histone acetylation and induce chromatin decompaction (intrinsically increasing NE tension) in different cells and also in HSCs^31–33^. RhoA activity levels were sharply upregulated in HSCs treated with 5mM NaB, in parallel with the increase in NE tension, as shown by the increase in the nuclear diameter and P-cPLA2 translocation to the NE (**Figure 1E-G** and **Video S5-6**)

To further support that P-cPLA2 translocation to the NE can be triggered by changes in NE tension^26^, we used hypotonic shock which was previously used to increase nuclear volume^22,25,34^ and NE tension and we quantified NE P-cPLA2 and RhoA activation. Consistently, the data shows an increase in NE P-cPLA2 and RhoA-GTP levels in HSCs after hypotonic osmotic shock (**Figure S1I**).

Overall, the data demonstrates that intrinsic changes in the NE tension activate RhoA in HSCs. However, RhoA activity might also be triggered by the extracellular matrix (niche or extrinsic activation). To explore this possibility, we focused on tissue stiffness, a well-described mechanical stimulus inducing RhoA activity in different cells and tissues^35,36^. Since in adult mammals HSCs reside in the BM, we first measured the stiffness of this compartment, which is a semi-solid tissue with viscoelastic properties and a quite heterogeneous mechanical behavior^37^. To this end, we used a Nanoindenter device equipped with a small and sensitive cantilever tip. We designed a matrix of several points to cover the whole area of the femoral BM tissue and obtain a map of the stiffness of the murine BM cavity (**Figure S1J**). In agreement with previous observations^38^, within the BM we detected areas with different levels of stiffness, ranging from 0.5 to 10kPa (on average 4kPa) in the inner marrow to a range of 5 to 50 kPa (on average 12kPa) at the endosteum (**Figure S1K**). Based on these results, we prepared polyacrylamide hydrogels functionalized with fibronectin (FN) to mimic *in vitro* the lowest (at the inner marrow; ∼1kPa) and the highest (endosteum; ∼40kPa) stiffness values that we detected in our murine BM samples. We then isolated HSCs and cultured them overnight on the functionalized hydrogels with different stiffness values. HSCs were then recovered and stained for RhoA-GTP and DAPI and used for a colony-forming unit (CFU) assay (**Figure S1L**).

Interestingly, RhoA-GTP levels did not changed in HSCs cultured on the stiff (40kPa) hydrogels compared to those cultured on the 1kPa hydrogels, contrary to what expected for RhoA being activated by an increased stiffness of the substrate^39^ (**Figure 1H**). As for the CFU assay, we did not detect differences in colony number, but the number of c-kit+ cells (hematopoietic progenitors) and the total cell number were reduced in the 40kPa-stiff hydrogels with no differences in the number of myeloid cells (Gr1+ and Mac1+ cells) (**Figure 1I-K** and **Figure S1M**). Intriguingly, RhoAKO HSCs behaved as their wild-type control for all the measured parameters (**Figure 1I-K** and **Figure S1M**).

Therefore, our data reveals that in HSCs RhoA is dispensable in the response to changes in extracellular stiffness, while RhoA is necessary to survive intrinsic changes in NE tension. In HSCs RhoA activity sharply increases after NE tension raise induced by confinement, osmotic shock and chromatin decompaction after NaB treatment (**Figure 1L**).

### RhoA activity is increased in aged HSCs

Aging alters the biomechanical properties of tissues and cells^40^. Since increased tissue stiffening upon aging has been reported to induce an increase in RhoA activity levels for example in the hair follicle^41^, we wondered if RhoA activity was altered in aged haematopoietic stem and progenitor cells (HSPCs) and whether it was correlated with increased stiffness of the aged BM^42^. We profiled the stiffness of the aged BM by performing indentation experiments using the same NanoIndenter device as reported above (**Figure S1J**). We collected several measurements alongside a matrix to cover the overall BM cavity of femurs and tibiae comparing in parallel young (8 to 20 weeks old) and aged C57Bl6 (>80 weeks old) mice. Strikingly, the indentation measurements showed an overall decreased bone marrow stiffness in aged samples compared to young, resulting in a homogenously low stiffness along the transversal section of the bone marrow (**Figure S2A-B**). To investigate whether the decreased BM stiffness was associated to changes in RhoA activity, we performed western blot and pull-down assays on HSPCs isolated from young and aged mice. We detected a significant upregulation of RhoA activity levels in aged BM cells, that could be targeted and significantly reduced by treatment with a selective RhoA inhibitor (Rhosin, here referred to as Ri)^43^ (**Figure S2C**). Further, we measured RhoA-GTP levels in sorted HSCs by immunofluorescence. To this end, HSCs were harvested from aged mice and incubated with or without Ri. Young HSCs were sorted alongside as control (**Figure 2A** and **Figure S2D**). Aged HSCs showed a dramatic upregulation of RhoA activity and treatment with 100µM Ri significantly decreased levels of RhoA-GTP in aged HSCs (**Figure 2B**), consistent with the western blot/pulldown results on HSPCs (**Figure S2C**). Therefore, in agreement with our conclusions based on the hydrogel experiments (**Figure 1I-K**), RhoA activity appears not to be associated to the extracellular stiffness and, while the aged BM stiffness decreases, RhoA activity increases in HSCs upon aging, supporting an intrinsic activation of RhoA in HSCs upon aging.

**Figure 2.**
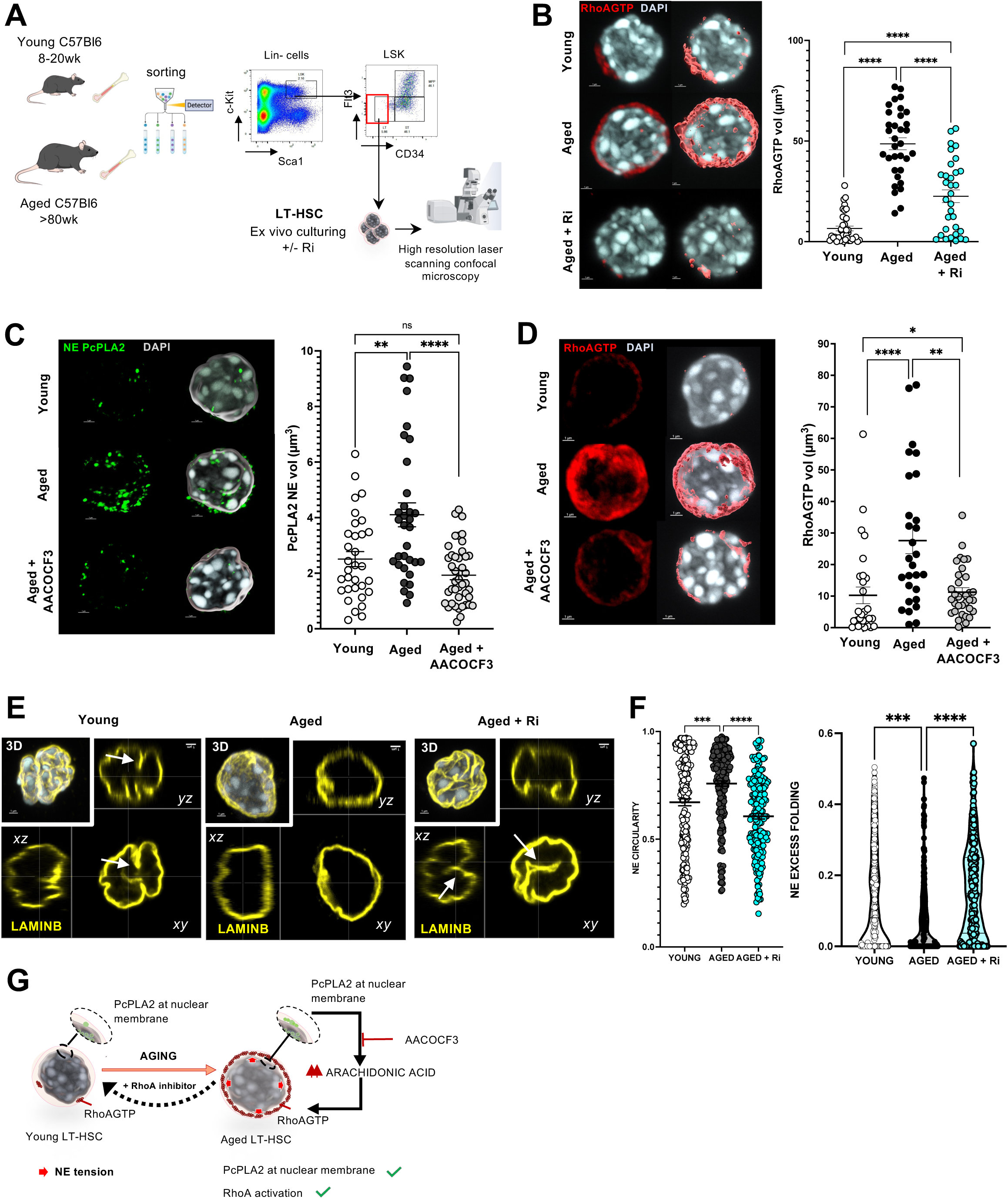
NE tension and RhoA activity are increased in aged HSCs. **A,** Representative HSC gating strategy and experimental set-up for *in vitro* culturing experiments with and without treatment. **B,** Representative images of 3D confocal reconstruction of young, aged and aged+Ri HSC stained with anti-RhoAGTP antibody (red) and DAPI (grey). Graph on the left shows RhoAGTP volume based on intensity of the signal. n=3, Mann-Whitney two-tailed test ****p<0.0001. Graph on the right shows HSC maximum diameter in *xy* for each condition. *p<0.05; **p<0.01 **C,** Representative images of 3D confocal reconstruction and analysis of young, aged and aged treated with AACOCF3 HSCs stained with anti-PcPLA2 showing compartmentalized PcPLA2 at the NE after segmentation. NE PcPLA2 is shown in green and DAPI in gray. The graph shows measurements of percentage volume of PcPLA2 at NE signal normalized against total volume of PcPLA2. n=3, Mann-Whitney two-tailed test ***p<0.001. **D,** Representative images of 3D confocal reconstruction of young, aged and aged+ AACOCF3 treated HSC stained with anti-RhoAGTP antibody (red) and DAPI (gray). The graph shows volume of RhoAGTP signal. n=3, Mann-Whitney two-tailed test *p<0.05; **p<0.01; ****p<0.0001. **E,** Representative images of 3D reconstruction and sections at the three axis *xy, xz* and *yz* of young, aged and aged Ri treated LT-HSC stained with LaminB in yellow and DAPI in gray. Arrows in white point to NE wrinkles. **F,** Nucleus wrinkling analysis. Nuclear envelope (NE) circularity of the nucleus (4𝜋𝐴⁄𝑃^2^, with 𝐴 the area of the nucleus and 𝑃 the total perimeter of the nucleus) for aged HSCs cells (n=29), aged HSCs treated with RhoA inhibitor (n=32) and young HSCs cells (n=30) from 3 independent experiments. Kruskal Wallis test for multiple comparisons (****p<0.0001 and ***p<0.001). Quantification of NE excess folding parameter (1 − 𝑝⁄𝑃, with 𝑝 the outline of the nucleus and 𝑃 the total perimeter of the nucleus) for aged HSCs (n=29), aged HSCs treated with RhoA inhibitor (n=32) and young HSCs cells (n=30) from 3 independent experiments. Kruskal Wallis test for multiple comparisons (****p<0.0001 and ***p<0.001). **G.** Cartoon scheme summarizing the findings.

Since we reported above that changes in NE tension can intrinsically activate RhoA in HSCs, we hypothesized that RhoA-GTP levels are higher in aged HSCs because their nucleus might be experiencing higher nuclear tension. Consistent with this hypothesis, we detected a significant increase of NE P-cPLA2 in aged stem cells (**Figure 2C** and **Figure S2E**). By treatment with the cPLA2 inhibitor (AACOCF3), NE translocation of P-cPLA2 is clearly reduced across all measurement metrics (**Figure 2C** and **Figure S2E**). Importantly, levels of RhoA-GTP in aged cells were sharply reduced to levels similar to young HSCs after treatment with AACOCF3, supporting that changes in NE tension are necessary to activate RhoA in aged HSCs (**Figure 2D**).

To further investigate the increased NE tension in aged stem cells, we performed additional experiments to measure the wrinkling of the NE, a structural feature of nuclear architecture that has previously been used as a measure of NE tension^22,25,26,44,45^. LaminB staining clearly shows increased NE circularity and reduced NE excess folding in aged HSCs compared to young (**Figure 2E-F**). This data strongly supports the results based on NE P-cPLA2 increase in aged stem cells and therefore that the NE tension in aged HSCs is higher compared to young stem cells (**Figure 2E-F**). Strikingly, RhoA inhibition sharply decreases NE circularity and NE excess folding in aged HSCs (**Figure 2E-F**).

To corroborate further these observations, we quantified also the nuclear import of the mechanosensitive transcription factor TAZ, which is known to accumulate in the nucleus with increasing NE tension^46,47^. In agreement, we measured increased nuclear translocation of TAZ after 8µm and 5µm HSC confinement (**Figure S2F**). As reported also previously^48^, we quantified higher level of nuclear TAZ in aged HSCs compared to young, which is dependent on RhoA activity (**Figure S2F**). Altogether, the results support the functional connection between changes in NE tension and RhoA activity as a mechano-sensitive regulator of HSC ageing (**Figure 2G**).

### RhoA inhibition restores DAPI-Intense Regions in aged HSCs

To explore if changes in NE tension impact on chromatin of aged HSCs, we developed a computational approach leveraging image analysis algorithms to extract morphometric and fluorescence intensity features from 3D-confocal images of HSC nuclei stained with DAPI, which is a photostable fluorescent DNA dye and its fluorescence intensity has been used in several applications to quantify DNA amount and chromatin condensation in intact nuclei^49,50^. First, consistent with increased NE tension, the results show that aged HSCs display a significant increase in nuclear volume, nuclear diameter, surface area, perimeter of the largest Z slide and DAPI-Intense Regions (DIRs) volume, among other related features, compared to young HSCs (**Figure 3A**). To note, HSC confinement (8µm) induced a similar larger increase in the same morphometric features, supporting that these alterations of nuclear volume, size and shape in aged HSCs are compatible with increased NE tension (**Figure 3A**)^19,25,26,51^.

**Figure 3.**
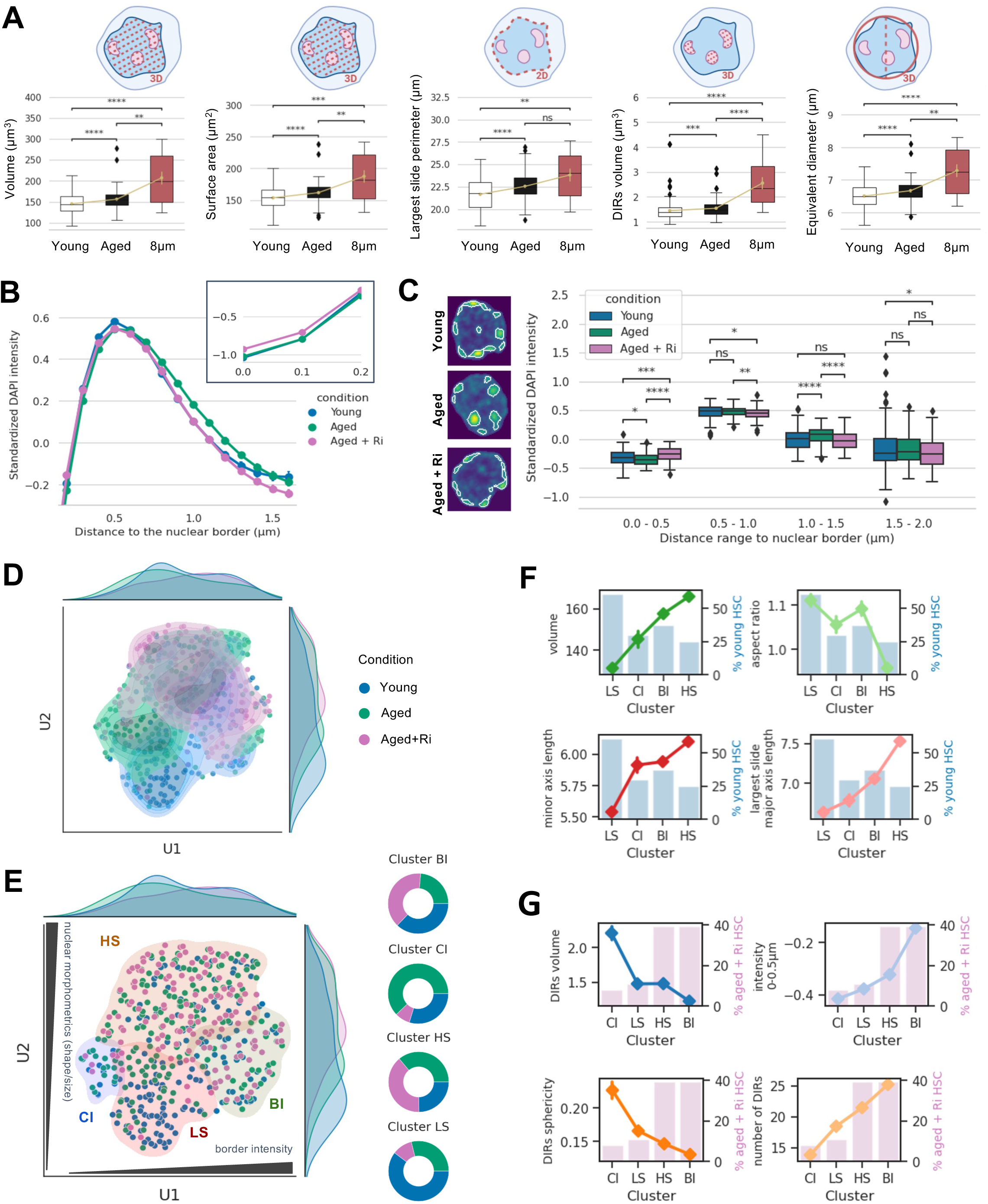
Drivers of aging identified by individual image-derived morphometric and DAPI intensity features. **A,** Boxplot displaying differences in selected feature distributions among HSCs young, aged, and young confined to 8 μm. Mean values are represented by a yellow dot on top of each distribution, with a yellow line aiding to visualize trends among conditions. Statistical significance was determined using the Mann-Whitney test. **B.** Line plot depicting the normalized DAPI intensity of nuclei obtained from young, aged, and aged + Ri conditions as a function of 3D iso-distant intervals of 0.1μm for up to 1.5μm from the nuclear border. A magnified view of the intensity within the initial 0.2 μm from the nuclear border is provided on the top right. **C.** Representative images of young, aged, and aged + Ri nuclear intensity images are shown, with DIRs highlighted by a white contour. Boxplots showing the standardized DAPI intensity of young, aged, and aged + Ri nuclei as a function of distance in 0.5 μm intervals. Statistical significance was determined using the Mann-Whitney test. **D.** Scatterplot displaying the UMAP embedding of all nuclei images, with marginal distributions for both components, colored by condition. The estimated kernel density helps to visualize high-density areas for each condition. **E.** UMAP embedding with background areas colored by clusters identified using the K-Means clustering algorithm (BI: Border Intensity, CI: Central Intensity, HS: High Size, LS: Low Size). Donut charts depicting the proportion of each condition per cluster are included on top. Marginal distributions for both UMAP components are also displayed. **F**. Lineplots showing the change in average values for selected morphometric-related features across clusters, ordered from the cluster with highest to lowest frequency of young HSCs. **G.** Lineplots showing the change in average values for selected intensity-related features across clusters, ordered from the cluster with lowest to highest frequency of aged+Ri HSCs.

Next, since RhoA inhibition sharply decreases NE circularity and NE excess folding in aged HSCs (**Figure 2E-F**), we asked whether decreasing RhoA activity might feedback to the nucleus inducing any change in morphometric and fluorescence intensity features of aged HSC nuclei. Surprisingly, the data revealed differences mainly in the pattern of DAPI intensity, that we extracted by quantifying fluorescence intensities along 3D iso-distant intervals from the nuclear border (**Figure S3A** and **Figure 3B)**. The pattern of DAPI intensities showed lower values near the nuclear border for aged HSCs compared to young and in aged HSCs DIRs localize relatively far from the nuclear border towards the central part of the nucleus (**Figure 3C**). Most aged+Ri HSCs nuclei displayed a large proportion of DIRs near the NE, similar to young HSCs (**Figure 3C**). In addition, other DAPI intensity features were different between young and aged HSCs and were restored to a youthful level after RhoA inhibition, namely DIRs height, DIRs major axis length, DIR distance to the border and number of DIRs (**Figure S3B**).

Next, as we calculated multiple morphometric and intensity features, we proceeded to conduct dimensionality reduction analyses on our feature set to explore similarity patterns within our HSC nucleus images (**Figure S3C-D**). First, we performed feature engineering and clustering analysis on all available nucleus images to uncover potential mechanisms underlying nuclear remodeling in young, aged and aged Ri-treated HSCs. The extracted imaging features are categorized into three groups: (*i*) whole nucleus level features, (*ii*) DIRs level features, and (*iii*) features computed from the largest 2D *z* slide in the *xy* plane. A comprehensive summary of these features is provided in **Table S1**. To address the high dimensionality and notable correlation of our feature space, we employed the non-linear dimensionality reduction technique Uniform Manifold Approximation and Projection^52^ (UMAP). Feature selection involved identifying statistically significant features (Mann Whitney U-test p-value < 0.05) among young and aged HSCs, and among aged and aged Ri-treated HSCs (**Figure S3C-D**). These significant features were then combined and filtered to remove those exhibiting high Pearson correlation coefficient (absolute value of correlation > 80%), resulting in a final set of 20 features (**Figure S3E**). The UMAP revealed that young, aged and aged+Ri HSCs exhibit overlapping yet distinct distributions, indicating underlying differences in their chromatin properties (**Figure 3D**). By grouping the nucleus data points using the K-Means clustering algorithm on the original feature space, we identified four distinct clusters that differ in morphometrics and intensity characteristics (**Figure 3E** and **Figure S3F-G**). Analysis of individual features’ impact over the UMAP representation revealed that morphometric-related features polarize the embedding vertically, whereas intensity and DIRs-related features polarize the embedding horizontally (**Figure 3E** and **Figure S3H**). Subsequently, by assessing the most representative features and biological populations, we annotated the clusters as Low Size (LS), High Size (HS), Border Intensity (BI) and Central Intensity (CI) (**Figure 3E**). Feature importance for each cluster was evaluated by measuring feature statistical significance and fold change per cluster (**Figure S3I**). Cluster BI includes mostly young and aged+Ri HSCs exhibiting high intensity near the NE and lower intensity in the nuclear center, along with an increased number of small DIRs closer to the border, which tend to be less spherical compared to nuclei in other clusters (**Figure 3E-G** and **Figure S3H-I**). Cluster CI predominantly consists of aged nuclei, with a reduced number of aged Ri-treated nuclei. These nuclei present with larger DIRs located farther from the NE and are notably spherical, with decreased intensity near the nuclear border (**Figure 3E-G** and **Figure S3H-I**). Cluster LS mainly contains young nuclei characterized by smaller size and lower mean intensity of DIRs, with surprisingly not much accentuated intensity values near the border (**Figure 3E-G** and **Figure S3H-I**). Cluster HS contains a mix of biological conditions, with nuclei characterized by larger size, DIRs positioned away from the nuclear border, and a decreased aspect ratio indicating these nuclei are wider than they are tall (**Figure 3E-G** and **Figure S3H-I**). By plotting feature trajectories across clusters, morphometric features progressively decrease or increase with decreasing frequencies of young HSCs within the clusters spanning from LS to HS clusters (**Figure 3F**). Interestingly, intensity features, and DIR-related features progressively decrease or increase with increasing frequencies of aged+Ri HSCs from CI to BI clusters (**Figure 3G**). Therefore, aged Ri-treated nuclei appear to share some morphometric similarities with aged nuclei yet exhibit intensity and DIR-related characteristics notably similar to those of young HSCs.

Overall, our computational approach suggests that DAPI imaging features elucidate chromatin differences in stem cells revealing a Ri-associated nuclear remodeling, which is mainly linked to changes in DAPI intensity and DIR volume, number and localization.

### RhoA inhibition restores H3K9me2 at heterochromatin

Alterations in the mechanobiology of the cell have been associated with aging-dependent changes in chromatin architecture^40,53^ and several epigenetic alterations characterize intrinsic HSC aging^54–56^, among which it has been previously reported a general loss of heterochromatin^3,57,58^. Intrigued by the observation that in aged HSCs treated with Ri some nuclear DAPI intensity and DIR-related features were significantly reverted to the level found in young HSCs (**Figure 3C** and **Figure S3B**) and are associated to the nuclear remodeling induced by Ri (**Figure 3G**), we focused on heterochromatin because especially DIRs are related to the most condensed portion of chromatin. Therefore, we analyzed the levels and distribution of H3K9me2, a known heterochromatin histone mark which is altered in aged HSCs^3,33^. By 3D-IF analyses, we measure a significant increase of H3K9me2 levels in aged+Ri HSCs compared to aged controls, together with a partial re-localization at the nuclear border like in young HSCs (**Figure 4A**). Interestingly, treatment with UNC0631, a selective inhibitor of the histone methyltransferase G9a specific to H3K9me2^59^, blunts completely the effect of Ri on H3K9me2 in aged HSCs (**Figure 4A**). This data suggests the requirement of G9a to increase levels of H3K9me2 in aged+Ri HSCs. Moreover, it reveals that aged HSCs are not affected by UNC0631 treatment alone, most likely because of the very low expression of G9a in these cells in basal conditions, which is partially rescued by Ri treatment (**Figure S4A**). To causally explain the role of decreased H3K9me2 in HSCs, we transduced young hematopoietic stem and progenitor cells (HSPCs or LSKs, gated as Lin^-^c-Kit^+^Sca-1^+^) with a retroviral vector, codifying for a histone variant in which the lysine of H3K9 is replaced by an arginine (H3R9) (**Figure 4B** and **Figure S4B**). The arginine in position 9 on H3 cannot be methylated, enforcing H3K9me reduction in HSCs. We functionally validated the strategy and the H3R9 incorporation by transplanting transduced LSKs into lethally irradiated recipient mice. We measured a significant reduction of H3K9me in H3R9 myeloid progenitors (MPs) compared to control vector transduced H3K9 MPs (**Figure 4C**). Young H3R9 HSCs isolated from transplanted mice present with higher RhoA-GTP levels and increased nuclear stretching compared to controls (**Figure 4D**). Moreover, H3R9 HSCs show a clear premature-aging phenotype upon transplantations *in vivo*, characterized by expansion of LSK and granulocyte-monocyte progenitors (GMP), reduced BM and peripheral blood (PB) regeneration, reduced B-lymphopoiesis and myeloid skewing (**Figure 4E-G**). Altogether, this data demonstrates that reduced methylation of H3K9 causes nuclear stretching, increasing RhoA activation and driving aging of HSCs. Importantly, RhoA inhibition restores H3K9 methylation levels in aged HSCs.

**Figure 4.**
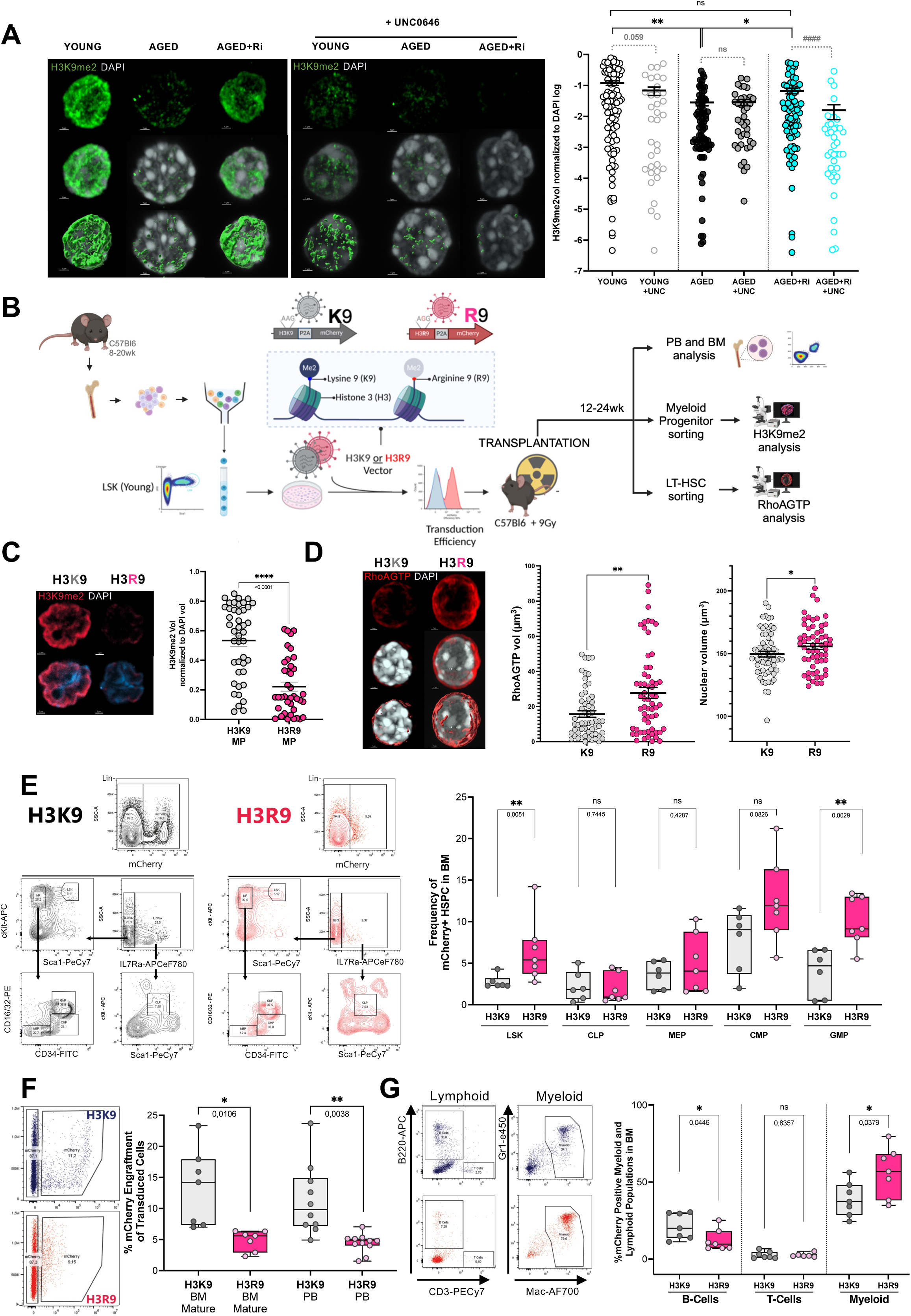
Reduced methylation of H3K9 increases RhoA activation and drives aging of HSCs. **A,** Representative images of 3D confocal reconstruction of young, aged and aged+Ri treated HSCs stained with anti-H3K9me2 antibody (green) and DAPI (grey). Representative images of 3D confocal reconstruction of young, aged and aged+Ri with UNC0631 treatment is shown. At least 10 cells were acquired per experiment per condition. n=4-8 Graph shows H3K9me2 signal volume normalized against DAPI signal volume. Mann-Whitney two-tailed, n=8, **p<0.01, *p<0.05, ####p<0.0001. **B.** Experimental set up for retroviral transduction and transplantation of young LSK overexpressing wild type H3K9 or mutant H3R9 with mCherry as a reporter. Transduced cells were transplanted into lethally irradiated mice. Bone marrow of transplanted mice was analyzed by flow cytometry to evaluate the regenerative capacity of transduced HSCs. mCherry^+^ Myeloid Progenitors (MP) were sorted to measure H3K9me2 levels. mCherry^+^ HSCs were sorted for the analysis of RhoA activation and nuclear stretching. **C.** Representative images of sorted MP transduced with H3K9 and H3R9 stained with anti-H3K9me2 antibody (magenta) and DAPI (blue). The graph shows H3K9me2 signal volume normalized against DAPI signal volume. Mann-Whitney two-tailed n=7 H3K9; n=6 H3R9, ****p<0.0001. **D.** Representative images of 3D confocal reconstruction of HSC overexpressing H3K9 or H3R9 isolated from transplanted mice (H3K9 n=7; H3R9 n=6). HSCs were stained with anti-RhoAGTP antibody (red) and DAPI (grey). Mann-Whitney, two-tailed statistics are shown for RhoAGTP quantification analysis **p<0.01. For nuclear volume (DAPI volume) unpaired t-test (one-tailed) was used *p<0.05. **E.** Gating strategy for the analysis of bone marrow progenitors after 12-20 weeks post transplantation. The analysis of the different populations is shown for mice transplanted with LSK transduced with either wild type H3K9 (n=7) or H3R9 (n=6). Mann-Whitney statistics are shown two-tailed **p<0.01. **F.** Engraftment of mCherry^+^-H3K9 or mCherry^+^-H3R9 HSCs in mice. Representative flow cytometry charts for bone marrow engraftment are shown. The box plots display engraftment of mCherry^+^-H3K9 or mCherry^+^-H3R9 HSCs in mice 12-20 weeks post-transplant in BM and peripheral blood. Unpaired t-test statistics is shown *p<0.05, **p<0.01. **G.** Gating strategy for the analysis of B cells (B220+), T cells (Cd3+) and myeloid cells (Gr1+ and Mac1+) within the BM after 12-24 weeks post transplantation. The analysis of the different populations is shown for mice transplanted with wild type H3K9 (n=7) transduced LSK and mutant H3R9 LSK (n=6). Mann-Whitney statistics are shown *p<0.05, **p<0.01.

### RhoA regulates chromatin accessibility at retrotransposons in aged HSCs

Previously, it has been proposed that the cell nucleus can directly respond to mechanical stress by inducing chromatin remodeling, altering polymerase and transcription factor accessibility and activity^60–62^, while increased chromatin accessibility as measured by ATAC-seq has been already characterized as an epigenetic alteration intrinsic to HSC aging^63,64^. Intrigued by possibility that RhoA inhibition might underscore a link between increased NE tension and increased chromatin accessibility in aged HSCs, we investigated further the chromatin remodeling associated with Ri treatment by performing ATAC-seq profiling of sorted young, aged, and aged+Ri HSCs (**Figure 5A**). Peak-calling identified a total of 57,289 accessible regions consistent between samples, most of them located in introns, intergenic regions, and gene promoters (**Figure S5A-B** and **Table S2**). In agreement with previously published data^63,64^, we detected an increase in open differentially accessible regions (DARs) with aging (2713 DARs open and 1103 DARs closed in aged compared to young HSCs; ∼8% FPR; **Figure 5B** and **Table S2**). After applying the Ri treatment to aged HSCs, 743 chromatin regions opened and 355 closed (∼8% FPR; **Figure 5B** and **Table S2**). Overall, 144 DARs were detected in both comparisons (aged *vs* young and aged+Ri *vs* aged), with 85.42% of them showing accessibility levels after the Ri treatment changing in the direction of the young levels (**Figure 5C** and **Figure S5C** and **Table S2**). Among the regions opening in aged HSCs treated with Ri, Gene ontology (GO) enrichment analysis revealed pathways related to cell migration, morphogenesis, adhesion, and chemotaxis (**Figure S5D** and **Table S2**). Interestingly, no GOs were significantly enriched among the DARs closing in aged HSCs+Ri and a high percentage of these closing DARs were located at retrotransposons (**Figure 5D lower right panel**), especially Long Terminal Repeats (LTRs) and Long INnterspersed Elements (LINE), like ERVL-MaLR, ERVK, ERV1, and L1 families (2-fold/∼1 log2FC higher percentage compared to the percentage in consensus peaks; FDR = 0.0018 for LTRs and FDR = 0.014 for LINEs in a one-proportion z-test; **Figure 5D-E** and **Table S2**)^65^. Notably, in aged HSCs we observe an opening of chromatin at retrotransposons, with a 1.65-fold (0.72 log2FC) and a 1.25-fold (0.33 log2FC) increase in the percentage of open DARs localizing in LTRs and LINEs, respectively (FDR = 3.2x10^-13^ for LTRs and FDR = 0.04998 for LINEs in a one-proportion z-test; **Figure 5D-E** and **Table S2**). Some of these DARs at LTRs and LINEs were overlapping with enhancers described previously^66^, while others were located in intronic and intergenic regions (**Figure 5E** and **Table S2**). Interestingly, retrotransposons and in particular LINE-1 have been suggested to directly contribute to aging of somatic cells and aging-related diseases^65,67–69^. To explore if the increase in chromatin accessibility at retrotransposons upon HSC aging is linked to increased NE tension, we performed ATAC-seq of young HSCs under 5μm confinement and compared their chromatin accessibility profiles to those of unconfined HSCs sorted in parallel from the same mice (**Figure 5F**). As a reference for DAR identification, we used the 42,632 consensus peaks identified between young and aged HSCs samples (**Table S3**). Data show 443 DARs opening and only 25 DARs closing in young HSCs under confinement (∼6.6% FPR; **Figure 5G-H** and **Figure S5E** and **Table S3**). Strikingly, while the DARs that are closing in confined HSCs are located at promoter-TSS and 5’UTR (1.67 log2FC over consensus peaks; FDR = 4.1x10^-6^ in a one-proportion z-test; **Figure 5I lower panel)**, the DARs that are opening are mainly at LTRs and LINEs (0.75 and 1.18 log2FC over consensus peaks, respectively; FDR = 0.008 for LTRs and FDR = 0.003 for LINEs in a one-proportion z-test; **Figure 5I upper panel**). Overall, the results support that the increased accessibility at REs observed in the aged HSCs and reverted (closed) by Ri treatment are located at LTR and LINE, which are the same type of genomic regions that are open by increasing NE tension by mechanical confinement. Next, we investigated transcriptional changes by bulk RNA-seq analysis of young, aged and aged+Ri HSCs. We identified 38 genes upregulated and 27 downregulated in aged HSCs after treatment with Ri (∼5% FPR; **Figure S5F-H** and **Table S4**). Consistently with the ATAC-seq data, after Ri treatment we detected a downregulation in the transcription of several retrotransposons subfamilies that are upregulated in aged HSCs, mainly LTRs but also some LINEs and DNA transposon subfamilies, like L1 (L1ME1 and L1ME3A), ERV1 (MER110-int) and hAT-Blackjack (MER63C) (∼8% FPR; **Figure 5J** and **Figure S5I** and **Table S4**). Gene Set Enrichment Analysis (GSEA) revealed enrichment for several GOs related to inflammation, innate immune response activation and interferon response among the genes downregulated after Ri treatment, again consistent with a downregulation of retrotransposons^65,68,70^ (FDR < 0.05; **Figure 5K-L** and **Figure S5J** and **Table S4**). Using GSEA, we also detected a decrease in the Interferome.org gene signature^71^ and of interferon-stimulated genes (ISG)^72^ in aged+Ri HSCs (**Figure 5M-N** and **Figure S5K-L**). Moreover, we measured a significantly negative enrichment score in aged+Ri *vs* aged samples for the HSC aging signature defined by Svendsen et al.^73^, while the same signature was clearly positively enriched in the aged samples compared to the young ones (**Figure 5O** and **Figure S5M**).

**Figure 5.**
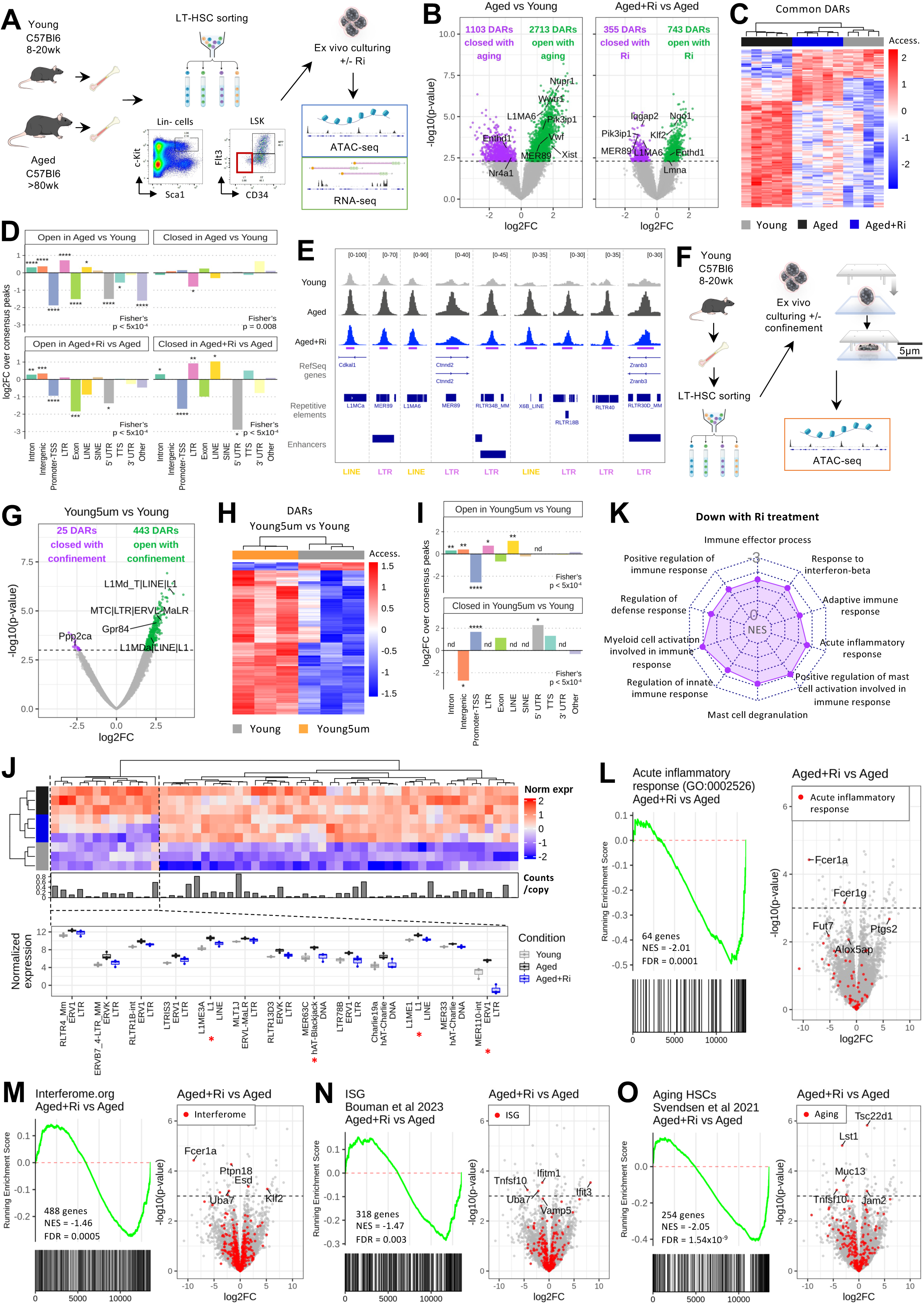
RhoA regulates chromatin accessibility at repetitive elements in aged HSCs. **A**, Representative HSC gating strategy and experimental strategy for ATAC-seq and RNA-seq of Ri treated aged HSCs. **B**, Volcano plot of DARs in aged *vs* young and aged+Ri *vs* aged (p<0.005). The annotation of selected DARs is shown. **C**, Heatmap of the normalized accessibility in the 144 common DARs in aged *vs* young and aged+Ri *vs* aged. **D**, Logarithmic fold change (log2FC) of the proportion of each genomic region type among the different sets of DARs over the proportions among the consensus peaks. Fisher’s exact test p-value is shown. One-proportion z-test for each genomic region is shown as *FDR<0.05, **FDR<0.005, ***FDR<0.0005, ****FDR<0.00005. **E**, Genome tracks for the normalized ATAC-seq read distribution in several LTRs and LINEs. Location of DARs is indicated with purple lines. **F**, Experimental strategy for ATAC-seq of young HSCs confined in 5μm. **G**, Volcano plot of DARs in young 5μm *vs* young (p<0.001). The annotation of selected DARs is shown. **H**, Heatmap of the normalized accessibility in the 468 DARs in young 5μm *vs* young. **I**, Log2FC of the proportion of each genomic region type among the different sets of DARs over the proportions among the consensus peaks. “nd” indicates that no regions of that type were detected as DARs. Statistics as in d. **J**, Heatmap of the normalized expression of the retrotransposon subfamilies upregulated with aging (p<0.005). Legend as in c. Boxplot shows the normalized expression per condition of the retrotransposon subfamiles downregulated with Ri (p<0.005). A red asterisk indicates the retrotransposon subfamilies that show significant differences in both comparisons. **K**, Radar chart showing the normalized enrichment scores (NES) for some significantly negatively enriched GOs with Ri (FDR<0.05). **L-O**, Enrichment plot and volcano plot for the acute inflammatory response GO term (L), the Interferome.org signature (M), the interferon-stimulated genes (N) and the aging signature (O) in aged+Ri *vs* aged. FDR = false discovery rate.

In summary, by ATAC-seq profiling, we detect changes in chromatin accessibility in aged Ri-treated HSCs that revealed reduction of open regions at LTRs and LINE, belonging mainly to ERVL-MaLR, ERVK, ERV1 and L1 families. ATAC-seq profiling of young 5µm-confined HSCs supports that the increased accessibility at retrotransposones observed in the aged HSCs and reverted by Ri treatment could be a consequence of increased nuclear stretching. Consistently, by bulk RNA-seq profiling of aged+Ri HSCs we detected a reduction in the transcription of LTRs and LINEs and a downregulation of the immune response, inflammatory response, interferon response and aging gene signatures compared to aged control HSCs.

### Inhibiting RhoA activity improves function of aged HSCs

Next, to further characterize the transcriptional changes in aged HSCs after Ri treatment, we focused on the few upregulated genes (**Figure S5G-H Table S4**) and, interestingly, we noticed three transcription factors (TF) belonging to the same family: *Klf4*, *Klf6* and *Klf2* (**Figure 6A** and **Table S4**). Noteworthy, the upregulation of Klfs is consistent with the opening of Klf4 motifs detected by ATAC-seq TF-binding motif analysis, which revealed enrichment in motifs for AP-1 and Klf4 (**Figure 6B** and **Table S2**). In hematopoietic cells, AP-1 can interact with chromatin remodelers to assist in the binding of other TFs^74^ and Klf4 has been previously shown to be important in cell reprogramming, blood formation and mechanosensing^75–80^. In addition, we identified an open DAR in aged+Ri HSCs in correspondence with a recently annotated Klf4 enhancer^81^ (**Figure 6C**) and we measured a clear increase in Klf4 protein in the nucleus of aged stem cells after Ri treatment (**Figure S6A**). Next, we analyzed the expression level of the genes targeted by a higher accessibility of Klf4-binding motifs and most of them increased their expression after Ri (**Figure S6B** and **Table S2**). GO enrichment analysis of these genes revealed enrichment for morphogenesis, differentiation, and actin polymerization (FDR < 0.05; **Figure 6D** and **Table S4**). Notably, actin polymerization (filamentous actin or F-actin) has been reported to restrict nuclear stretching^21,22^. Consistently, while in aged HSCs F-actin is decreased compared to young, F-actin levels increase upon Ri treatment similarly to young HSCs (**Figure 6E**).

**Figure 6.**
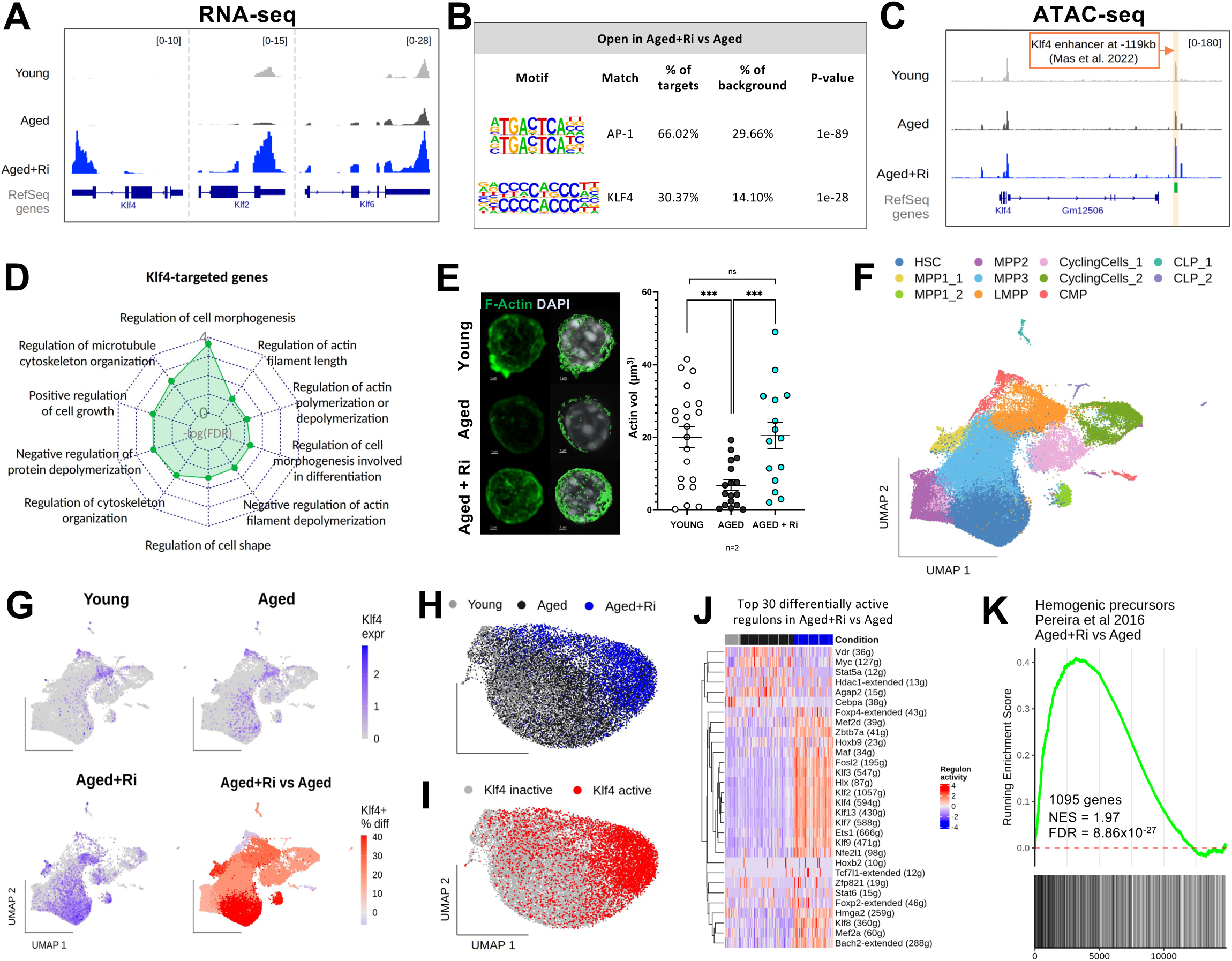
Inhibiting RhoA activity induces Klf4 expression and partial reprogramming in aged HSCs. **A**, Genome tracks for the normalized RNA-seq read distribution in Klf genes. **B**, Top 2 TF-binding motifs found among the DARs opening with Ri. **C**, Genome tracks for the normalized ATAC-seq read distribution in Klf4 gene and its enhancer in –119kb (orange shadow). A green line indicates the DAR opening with Ri. **D**, Radar chart showing the minus logarithmic FDR for some significantly enriched GOs among the Klf4-targeted genes (FDR < 0.05). **E**, Representative images of 3D confocal reconstruction of young, aged and aged+Ri LT-HSC stained with Phalloidin-488 (green) and DAPI (grey). The graph shows measurements of phalloidin (actin) signal volume, n=3. Mann-Whitney, two tailed ***p<0.001. **F**, Clustered and annotated UMAP for the integrated scRNA-seq data for young, aged and aged+Ri LSKs. HSC: hematopoietic stem cells; MPP: multipotent progenitors; LMPP: lymphoid multipotent progenitors; CMP: common myeloid progenitors; CLP: common lymphoid progenitors. **G**, Klf4 normalized expression in the UMAP for young, aged and aged+Ri. The lower right panel shows the difference in the percentage of Klf4+ cells in aged+Ri *vs* aged per cluster. **H**, UMAP based on the TF activity scores in HSCs colored by condition. **I**, Klf4 active cells (AUC > 0.128) in the UMAP. **J**, Heatmap of the activity scores for the top 30 TFs that are differentially active in aged+Ri *vs* aged (|log2FC| > 0.5 and FDR < 0.05). Color legend as in h. **K**, Enrichment plot for the hemogenic precursors signature in aged+Ri *vs* aged.

To gain further insights into the transcriptional rewiring of aged HSCs treated with Ri, we performed scRNA-seq of young, aged, and aged Ri-treated LSK cells. We obtained a total of 60,648 cells expressing 15,049 genes. Clustering of the integrated UMAP identified 11 clusters (**Figure 6F**) that were annotated as based on known marker gene expression and enrichment for several previously identified HSC and LSK signatures^82–84^ (**Figure 6C-F** and **Table S5)**. Compositional analysis of cell clusters showed the expected increase in the percentage of HSCs in all aged samples compared to the young ones (**Figure S6G** and **Table S5**). It also revealed an increase in the percentage of MPP3 with aging and its decrease with the Ri treatment (**Figure S6G** and **Table S5**). Interestingly, the scRNA-seq dataset showed that the increase of Klf4 expression in aged+Ri LSKs is more prominent in the HSC and the MPP1 clusters (**Figure 6G**). In detail, 43.58% of aged+Ri HSCs are expressing Klf4, while only 2.46% and 2.80% of young and aged HSCs, respectively, express this TF. Differential gene expression analysis between the three conditions in the HSC cluster confirms the upregulation of several genes of the Klf family after Ri treatment (|log2FC| > 1 and FDR < 0.05; **Figure S6H** and **Table S5**). To further analyze the activity of the different TFs and identify the network of regulated genes, we calculated TF activity scores in each single HSC using SCENIC and generated a UMAP based on TF activity scores (**Figure 6H**). Interestingly, the percentage of aged+Ri HSCs with active Klf4 is 89.52%, while it is negligible in young and aged HSCs (1.89% and 0.69% respectively) (**Figure 6I** and **Figure S6I**). Differential analysis of the activity scores in between conditions confirms the increased activity of several Klf TFs, as well as other TFs (|log2FC| > 0.5 and FDR < 0.05; **Figure 6J**). Since Klf4 is known for its role in cell reprogramming and dedifferentiation, we wondered if aged+Ri HSCs show a more dedifferentiated transcriptional state. Interestingly, GSEA reveals an enrichment of the hemogenic precursor signature defined by Pereira et al.^85^ in the aged+Ri HSCs compared to the aged controls (**Figure 6K**), supporting that Ri treatment might induce a partial transcriptional reprogramming of aged HSCs.

Since the scRNA-seq analysis indicates a partial reprogramming suggestive at the transcriptomic level of a possible functional improvement, we decided to assess the regenerative capacity of aged HSCs after Ri treatment *in vivo*. Previously, we correlated changes in cell epigenetic polarity of H4K16ac to function of HSCs^33,86–88^. 3D-IF staining of young, aged and aged+Ri HSCs clearly showed that Ri treatment restores H4K16ac polarity in aged stem cells (**Figure S7A**). Next, we tested in a non-competitive transplantation assay into young immunocompromised and *Kit^W-41J^* mutant mice (NBSGW) the regenerative capacity of aged Ri treated HSCs compared to young and aged control stem cells. We transplanted 200 aged HSCs treated overnight *ex vivo* with 100µM Ri, alongside control recipient mice transplanted with solvent treated young or aged HSC (**Figure 7A**). Donor mice were genetically tagged by constitutive expression in Rosa26 locus of a bright and stable red-fluorescent protein (tdRFP)^89^ and we measured donor-derived contribution in peripheral blood (PB) by detecting RFP^+^ cells at several time-points after transplantation. Notably, aged Ri-treated HSCs engrafted at the endpoint similarly to young HSCs, showing a significant increase in their peripheral blood (PB) regenerative capacity compared to aged control HSCs (**Figure 7B-C**). Remarkably, Ri treatment also significantly increased the B cell lymphoid differentiation potential of aged HSCs and decreased the contribution to the myeloid lineage 18 weeks after transplantation (**Figure 7B-D**). Ri treatment did not change the differentiation to the T cell compartment (**Figure 7B-D**). Engraftment in BM and HSC compartments was not significantly different in Ri treated recipients compared with either young or aged control recipients, showing a trend for both parameters to resemble young donor HSCs (**Figure S7B-C**). Overall, we conclude that inactivation of RhoA in aged HSCs functionally improves the regenerative capacity of old stem cells and their myeloid/B-lymphoid skewing *in vivo*.

**Figure 7.**
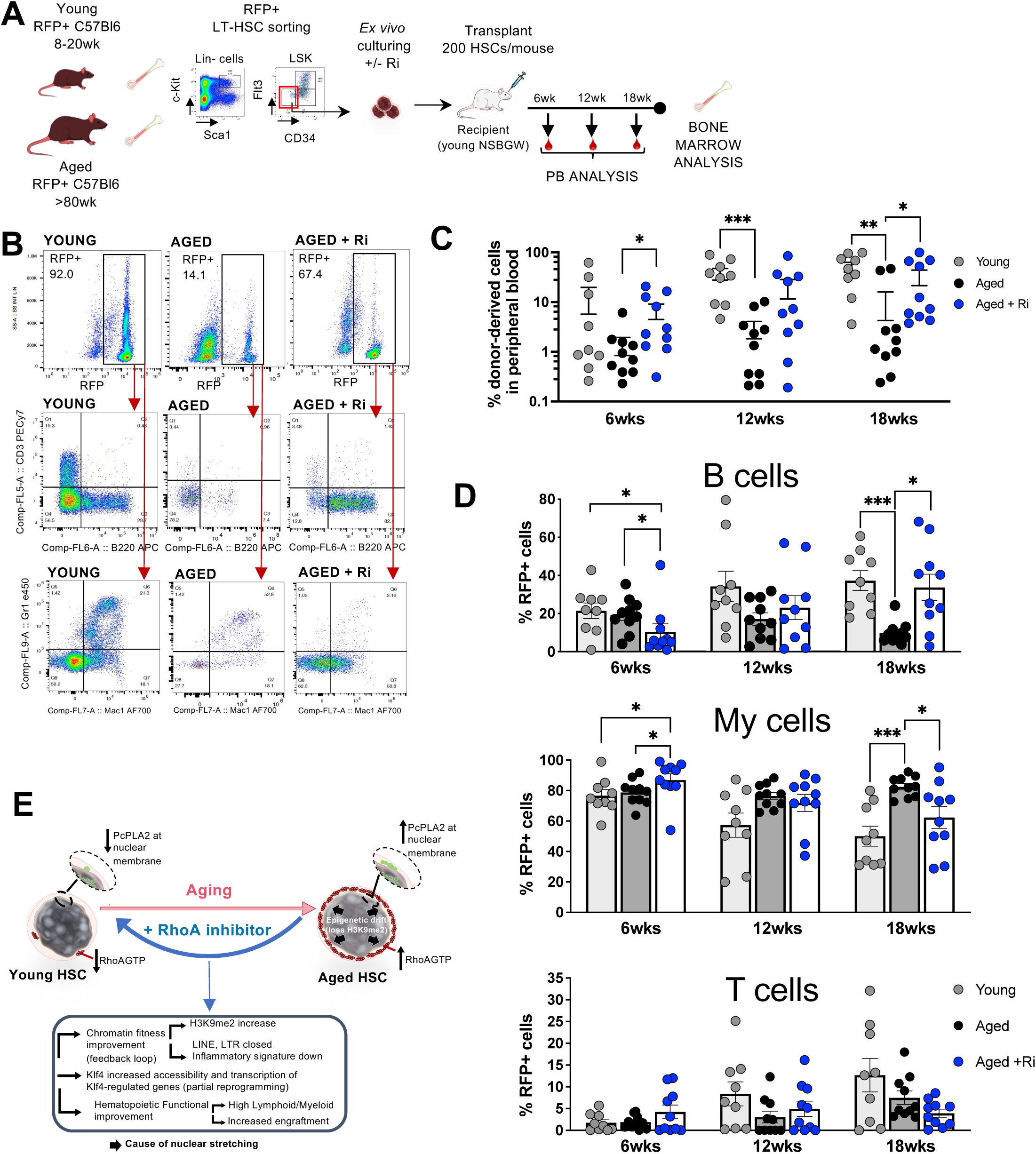
Inhibiting RhoA activity improves function of aged HSCs. **A**, Experimental strategy for transplantation of young, aged and aged+Ri RFP+ HSCs into NBSGW mice. **B**, Percentage of engraftment in PB along the course of the transplantation at 6, 12 and 18 weeks. 4 independent transplantation experiments were performed, and the initial number of recipient mice used per experiment was 3-4 per group. Young n=9; aged n=10; aged+Ri n=10. Mann-Whitney test, unpaired, two-tailed *p<0.05, **p<0.01, ***p<0.001. **C**, Representative flow cytometry gating strategies for RFP+ cells, lymphoid cells (CD3+ and B220+), and myeloid cells (Gr1+, Gr1+Mac1+ and Mac1+) in PB at 18 weeks. **D**, Graphs showing the percentage of donor derived B220+ cells, myeloid (Gr1+, Gr1+Mac1+ and Mac1+ cells) and CD3+ cells in PB at 6, 12 and 18 weeks after transplant. Young n=9; aged n=10; aged+Ri n=10.; Mann-Whitney test, unpaired two-tailed; \**p*<0.05, ***p<0.001. **E**, Cartoon scheme summarizing the main features rejuvenated by Ri treatment in aged HSCs.

## DISCUSSION

The regenerative potential of HSCs declines upon aging^90^. Moreover, aged HSCs are skewed toward myeloid differentiation, which contributes to immunosenescence and to the increased incidence of hematopoietic disorders in the elderly^1–4^. Previously, intrinsic epigenetic alterations have been associated with HSC aging, such as for example increased chromatin accessibility^3,33,54–56^. We also described that some of these epigenetic alterations depend on a reduction of LaminA/C expression in aged HSCs^33,63^, which is suggestive of a possible impairment of the mechanical properties of the nucleus. Supporting a novel emerging perspective that focuses on nuclear mechanoregulation^16,18,47,91,92^, here we investigate RhoA, a small GTPase critical for HSC cytokinesis and differentiation^20^ that has been involved also in mechanotransduction in different cell types^16^.

Our data shows that in HSCs RhoA is activated by increasing NE tension under cell confinement, by chromatin decompaction after NaB treatment, by nuclear swelling after hypotonic osmotic shock and after reduction of H3K9 methylation levels. To note, we identify the loss of methylation of H3K9 as a likely cause not only of RhoA over-activation and increased nuclear stretching in aged HSCs, but also of many phenotypes associated with functional aging of HSCs (reduced regenerative capacity, expansion of the primitive GMPs and LSKs populations and myeloid/lymphoid skewing in BM). Interestingly, G9a activity is required for Ri restoration of H3K9me2 levels in aged HSCs. Further work is necessary to clarify mechanistically how RhoA activity downregulation affects the activity of this histone methyltransferase. To note, the data reveals also that RhoA is necessary to survive cell confinement, which intrinsically induces RhoA activation. Differently from other cell types^39,41^, we show that in HSCs RhoA is not involved in transducing changes in extracellular stiffness, since RhoA is dispensable in the response to changes induced by culturing HSC on hydrogel with high stiffness.

Furthermore, our results reveal that NE tension is physiologically increased upon HSC aging and that NE tension is necessary to activate RhoA in aged HSCs. RhoA activity in aged HSCs can be targeted by a specific small molecule inhibitor^43,93^, which restores H3K9me2 levels and recovers the phenotypes of young HSC nuclei such as DAPI intensity, DIR volume, number and localization. This is further supported by our machine learning approach, which demonstrates that DAPI-imaging morphometric and intensity features can be used to explain differences between HSCs and to inform on the age and fitness of the stem cells.

Importantly, by ATAC-seq and RNA-seq we detect a downregulation in chromatin accessibility and transcription at LTRs and LINE and a decrease in inflammation, immune response, interferon responsive genes and aging signatures in aged Ri-treated HSCs. Moreover, after treatment of aged HSCs with Ri we measure an increased transcription of Klf4, an opening of Klf4-binding motifs and an increased activity of Klf4 TF. Genes targeted by Klf4 are enriched for pathways related to actin polymerization and the increased levels of F-actin are likely acting to restrict nuclear stretching in aged Ri-treated HSCs. Moreover, Klf4 targets also genes enriched for a signature of hemogenic precursors, compatible with a partial reprogramming of aged HSCs, supporting the functional improvement in the regenerative capacity and myeloid/lymphoid skewing of aged+Ri stem cells in transplantation assays (**Figure 7E**).

So far, several reports associated the aging process of different cell types with an epigenetic drift involving loos of heterochromatin and H3K9me and a dis-regulation of normally silenced and closed retrotransposons^65,67,68,94,95^. Here we report that an intrinsic nuclear mechanosignalling pathway dependent on RhoA can be pharmacologically targeted to revert these drifted epigenetic features, improving function of aged somatic stem cells (**Figure 7E**). To note, overactivation of RhoA has been previously reported to be associated with functional impairment of human HSCs, supporting possible translations of our findings^96^. Altogether our data sheds light on a new perspective of intrinsic nuclear mechanotransduction to control the aging-related epigenetic drift of somatic stem cells as a potential target for improving tissue homeostasis over time.

## Data availability statement

The source data underlying **Figure 1-6** and **Figure S1-6** is provided as a **Source Data file**. The source code for the microscopy image analyses showed on **Figure 3** and **Figure S3** is available at HYPERLINK “https://github.com/biomedical-data-science/hsc_rhoa”. Sequencing data underlining **Figure 4-5** and **Figure S4-5** and **Table S2-5** is deposited together with codes under the repository DOI https://doi.org/10.34810/data697. ATAC-seq, RNA-seq and scRNA-seq data are deposited at GEO (accession number GSE233989). Dilutions and catalogue numbers of all commercial antibodies are provided in the **Source Data file**.

## Acknowledgments & Funding Sources

We thank support from Dr. Laia Traveset Martinez, IDIBELL Innovation Unit. We acknowledge support from Dr. Mercè Marti Gaudes, head of Technical Facilities at IDIBELL together with José Andres Vaquero (IDIBELL FACS and Flow cytometry SCT), Antoni Ventura (IDIBELL Mouse Facility SCT) and Joan Repulles and Saioa Mendizuri (IDIBELL Bioimaging SCT). We thank Esther Castaño, Beatriz Barroso and Benjamin Torrejon (CCiT-UB, Bellvitge). We thank Giulia Lunazzi and the National Center for Genomic Analysis (CNAG, Barcelona) for the support with scRNA-seq experiments. We thank CERCA Program/Generalitat de Catalunya for institutional support. We thank Conxi Lazaro (LCAM laboratory, ICO-HUB) for supporting sequencing experiments. We acknowledge the funding sources: European Research Council (ERC) grant 101002453 (MCF), Spanish Ministry of Science, Innovation and University grants RYC2018-025979-I (MCF) and PGC2018-102049-B-I00 (MCF) and INPhINIT Incoming fellowship from “la Caixa’’ Foundation (ID 100010434) with code LCF/BQ/DI22/11940001 (PIP). VR acknowledges financial support from the Ministerio de Ciencia e Innovación through the Plan Nacional (PID2020-117011GB-I00) and funding from the European Union’s Horizon EIC-ESMEA Pathfinder program under grant agreement No 101046620.

## Declaration of interests

The findings presented in this study are covered under patent application number EP25382180.5, filed on 27/02/2025.

## Author contributions

Conceptualization: EM-R, MCF, PIP, BW

Methodology: SM-V, EM-R, MCF, PIP, FP, FA, JLC, BW, FM, LR, SW

Investigation: SM-V, EM-R, MCF, PIP, BW

Visualization: SM-V, EM-R, PIP, LM

Funding acquisition: MCF

Project administration: MCF

Supervision: YZ, VR, AR, PP, MCF

Writing – original draft: EM-R, SM-V, MCF, PIP

Writing – review & editing: AR, VR, YZ, PP, MCF

## Supplementary Figure Legends

**Figure S1| Related to Figure 1. A.** Confinement device scheme explaining the mounting strategy and how the cells are confined. **B.** Gating strategy for the isolation of LT-HSC from young BM cells from a representative experiment. **C.** Stripplot showing the increase of the excess of perimeter (EOP) upon confinement measured in 2D. **D.** Stripplot showing an increase in major nuclear axis length upon confinement at different heights. **E,** DAPI intensity profiles of representative confocal images from HSC under different levels of confinement as a function of the distance to the nuclear border. **F,** RhoA *locus* genetic maps of RhoA*wt* and RhoA*fl* mice and schematic result of *in vitro* Cre tamoxifen-induced deletion of RhoA exon3. Black arrowhead stands for the *flox* sequence location at the RhoA *locus*. PCR results are shown for the RhoA *floxed* (RAp1&RAp2) and RhoAKO (RAp1&RAp3) pair of primers performed on BM cells genomic DNA from Cre^+/-^ RhoA^wt/wt^ and Cre^+/-^RhoA^fl/fl^ mice after *in vitro* tamoxifen treatment and control. Sizes of expected PCR products are shown in each case. **G,** Representative confocal images, and their respective 3D reconstruction in LT-HSC after o/n treatment with tamoxifen for induction of RhoAKO. Cells were stained against RhoAGTP (red) and DAPI (gray). Graph shows the volume for RhoAGTP signal in wt and induced RhoAKO without confinement. Mann-Whitney, unpaired two-tailed is shown *p<0.05. **H.** Representative image of NE PcPLA2 (green) in 5µm confined HSC. Graph comparing levels of PcPLA2 at the NE in unconfined and 5µm confined cells. Mann-Whitney, unpaired two-tailed is shown ****p<0.0001, n=3. **I,** Representative image of NE PcPLA2 (green) and RhoA-GTP (red) in young HSC cultured in isotonic and 0.75X hypotonic conditions. DAPI is shown in grey. The graphs compare levels of NE PcPLA2 and RhoA-GTP in isotonic and hypotonic samples. Mann-Whitney, unpaired two-tailed is shown **p<0.01, *p<0.05. n=3. **J,** Schematic representation of the Chiaro NanoIndenter and design of bone measurement experiment. **K,** Graph showing the stiffness of the endosteum and inner marrow within femurs from young mice. A stiffness variation graph of a representative single femur is shown. Mann-Whitney, unpaired two-tailed ****p<0.0001. **L,** Schematic representation of the experimental design for the 2D culturing of LT-HSC on different stiffness hydrogels and later analysis by immunofluorescence (IF) and methocult culturing. **M,** Graph representing the number of cells obtained after 10 days on methocult from RhoAwt and RhoAKO LT-HSC cultured in different stiffness hydrogels. Mann-Whitney, unpaired two-tailed.

**Figure S2| Related to Figure 2. A,** Superposed stiffness variation graphs showing the measurements of representative femurs on a young and an aged mouse. **B,** Stiffness of the BM of femurs of young and aged mice at the endosteum and inner marrow. n=3, with a total of 69-80 measurements per condition, Mann-Whitney, two-tailed test *p<0.05; ****p<0.0001. **C,** Representative western blot of RhoAGTP pulled down from BM cells from young, aged and aged treated *in vitro* with Ri. Antibody against RhoA was used to detect pulled down RhoAGTP and total RhoA in whole protein extracts. Western blot of paired protein extraction samples against RhoA total and actin is shown as reference. n=3, Mann-Whitney test, unpaired one-tailed *p<0.05. **D,** Gating strategy for the isolation of LT-HSC from young and aged BM cells from representative experiments. **E.** Representative images of 3D confocal reconstruction and analysis of young, aged and aged treated with AACOCF3 HSCs stained with anti-PcPLA2. These images show total PcPLA2 (green) and the NE PcPLA2 signal (red) which overlaps completely with the total signal and appears in yellow (merged of red and green). Yellow arrows point to representative accumulation of NE PcPLA2 which can be more or less visible also depending on 3D rotation of the cells. Graphs are showing total PcPLA2 vol in µm^3^, percentage of NE PcPLA2 volume relative to total PcPLA2 volume and relative to DAPI volume. n=3 Mann-Whitney, two-tailed test *p<0.05; **p<0.01; ***p<0.001 ****p<0.0001. **F,** Representative images of 3D confocal reconstruction stained with anti-TAZ and its analysis of HSC from young, aged and aged treated with Ri along confined young cells at 8 µm and 5µm. n=3, Mann-Whitney, two-tailed test *p<0.05; **p<0.01; ***p<0.001 ****p<0.0001.

**Figure S3| Related to Figure 3. A.** Schematic representation of the analyses of DAPI intensity by distance to the nuclear border. **B.** Stripplot displaying differences in morphometric and DIRs-related feature distributions among young, aged, and aged Ri-treated nuclei. Mean values are represented by a yellow dot on top of each distribution, with a yellow line aiding to visualize trends among conditions. Statistical significance was determined using the Mann-Whitney test. **C.** Heatmap depicting the correlation matrix for the 11 significant features identified among young and aged nuclei, after discarding highly correlated features. Marginal dendrograms exhibit clusters of feature similarity. The accompanying UMAP embedding, fitted with the specified features, is shown on the right. **D.** Heatmap depicting the correlation matrix for the 11 significant features identified among aged and aged Ri-treated nuclei, after discarding highly correlated features. Marginal dendrograms exhibit clusters of feature similarity. The accompanying UMAP embedding, fitted with the specified features, is shown on the right. **E.** Heatmap depicting the correlation matrix for the 20 features combined from the previous pairwise comparisons (young and aged, aged and aged Ri-treated) after discarding highly correlated features. **F.** Silhouette plot showing the silhouette scores for K-Means clustering (K=4) for each nucleus in our dataset, with scores colored by cluster assignment. A red dashed vertical line indicates the overall silhouette score. **G.** Schematic representation of the multidimensional analysis pipeline. Features are extracted from whole DAPI intense microscopy images and used to construct a lower dimensional representation. Feature selection involved identifying statistically significant features among young and aged, and among aged and aged Ri-treated nuclei, followed by correlation-based filtering**. H.** Scatterplots illustrating the UMAP embedding constructed using nuclei from young, aged, and aged Ri-treated conditions, with color indicating variations in selected morphometric and intensity-related features. **I.** Volcano plots showing the most significant features among clusters. The x-axis represents the log2 feature fold change of the indicated cluster against the remaining clusters, while the y-axis indicates the significance of this change. Dot color intensity corresponds to significance, and the red horizontal line marks a significance threshold at a p-value of 0.005.

**Figure S4| A**, Representative images of 3D confocal reconstruction of young, aged and aged+Ri treated HSCs stained with G9a antibody (red) and DAPI (grey). Mann-Whitney two-tailed, **p<0.01, *p<0.05. **B**. Flow cytometric analysis of transduced LSK pre-transplantation, representing 3 independent batches of transplantation. Mann-Whitney test, unpaired two-tailed, p=0.700.

**Figure S5| Related to Figure 4. A**, Proportions of the consensus peaks annotations in a donut chart. **B**, PCA plot for the ATAC-seq normalized read counts in the consensus peaks for the Ri-treatment experiment. **C**, Venn diagram for the number of common DARs in aged *vs* young and aged+Ri *vs* aged (p<0.005). **D**, Radar chart showing the minus logarithmic FDR for the significantly enriched GOs among the genes annotated to the DARs opening with Ri (FDR<0.1). **E**, PCA plot for the ATAC-seq normalized read counts in the aged *vs* young DARs for the young confined HSCs experiment. **F**, PCA plot for the RNA-seq normalized read counts in genes. Legend as in d. **G**, Volcano plots of differentially expressed (DE) genes in aged *vs* young and aged+Ri *vs* aged (p<0.001). Selected genes are shown. **H**, Venn diagram for the number of common DE genes in aged *vs* young and aged+Ri *vs* aged (p<0.001). **I**, Venn diagram for the number of common DE retrotranspon subfamilies in aged *vs* young and aged+Ri *vs* aged (p<0.005). **J-M**, Enrichment plot and volcano plot for the acute inflammatory response GO term (J), the Interferome.org signature (K), the interferon-stimulated genes (L) and the aging signature (M) in aged+Ri *vs* aged. NES = normalized enrichment score. FDR = false discovery rate. Number of genes used for the analyses is also indicated.

**Figure S6| A**, Representative images of 3D confocal reconstruction of young, aged and aged+Ri treated HSCs stained with Klf4 antibody (red) and DAPI (grey). Mann-Whitney two-tailed, *p<0.05. Experimental mice per arm n=2. **B**, t-statistic for the Klf4-targeted genes in the aged+Ri *vs* aged comparison in ATAC-seq and RNA-seq. The number of genes with positive (green) and negative (purple) RNA-seq t-statistic is shown. Top upregulated genes are shown. **C**, Main marker genes used for cluster annotation. **D**, Cell scores for the HSC signature (MolO) defined by Wilson et al. **E-F**, Cell assignments for the low/high-output signature by Rodriguez-Fraticelli et al. (E) and the dormant/active signature by Cabezas-Wallscheid et al. (F). **G**, Percentage of cells in each cluster per sample. scCODA’s statistically credible changes are shown. *Effects Final Parameter (EFP)<0.5; **EFP<1; ***EFP<1.5. **H**, Log2FC against the average normalized expression for the DE analysis in aged *vs* young and aged+Ri *vs* aged in the HSC cluster (|log2FC| > 1 and FDR < 0.05). Selected genes are shown. **I**, Ctcf and Hoxa10 active cells (AUC > 0.007 and 0.01, respectively) in the UMAP.

**Figure S7| A**, Representative images of 3D confocal reconstruction of young, aged and aged+Ri HSCs stained with anti-H4K16ac antibody (green), RhoAGTP (red), and DAPI (gray) as reference. Dotted line indicates symmetry axes. The graph shows quantification of polar cell number for H4K16ac. N=3-5. 50 cells were counted per sample, per experiment, one-way Anova, *p<0.05; **p<0.01. **B**, Percentage of engraftment in bone marrow at 20wk post-transplantation, n=4 Mann-Whitney test, unpaired two-tailed. **C**, Percentage of HSC derived from transplantation 20wk after transplantation, n=4 Mann-Whitney test, unpaired two-tailed.

**Table S1 | Morphometric and intensity features extracted from DAPI microscope images used for the multivariate feature analysis**

**Table S2 | Data for the ATAC-seq experiment on young, aged and aged+Ri HSCs.** Information on the samples, the consensus peaks, the DARs, the GO enrichment, the TF motif analysis, and the Klf4-targeted genes.

**Table S3 | Data for the ATAC-seq experiment on young and young 5μm confined HSCs.** Information on the samples, the consensus peaks, and the DARs.

**Table S4 | Data for the RNA-seq experiment on young, aged and aged+Ri HSCs.** Information on the samples, the DE genes, the DE retrotransposons, the GSEA, and the GO enrichment of upregulated Klf4-targeted genes.

**Table S5 | Data for the scRNA-seq experiment on young, aged and aged+Ri LSKs.** Information on the samples, the cluster markers, the compositional analysis, the DE genes, and the differentially active regulons.

**Video S1 | Related to Figure 1**. Representative 3D immunofluorescence reconstructions of an uncofined HSC. The nucleus is stained by DAPI (gray). RhoAGTP was stained with an RhoAGTP antibody, and is shown in red. Scale bars=1 µm.

**Video S2 | Related to Figure 1**. Representative 3D immunofluorescence reconstructions of 8µm-confined HSC. The nucleus is stained by DAPI (gray). RhoAGTP was stained with an RhoAGTP antibody, and is shown in red. Scale bars=1 µm.

**Video S3 | Related to Figure 1**. Representative 3D immunofluorescence reconstructions of 5µm-confined HSC. The nucleus is stained by DAPI (gray). RhoAGTP was stained with an RhoAGTP antibody, and is shown in red. Scale bars=1 µm.

**Video S4 | Related to Figure 1**. Representative 3D immunofluorescence reconstructions of 3µm-confined HSC. The nucleus is stained by DAPI (gray). RhoAGTP was stained with an RhoAGTP antibody and is shown in red. Scale bars=1 µm.

**Video S5-6 | Related to Figure 1**. Representative 3D confocal reconstructions of HSCs showing phospho-cPLA2 (PcPLA2) (light pink) and DAPI (gray) in young control (Movie5) and young NaB (Movie6) treated HSCs. By using image analysis software Imaris we have compartmentalized PcPLA2 signal within the nucleus and at the nuclear envelope (NE). We have assigned light pink to NE PcPLA2 and magenta to nuclear PcPLA2 after compartmentalization. Surfaces have been created for better visualization. Scale bar=1µm

## Methods

### Reagents

A list of reagents, chemicals, commercial kits and antibodies is provided as **Source Data file**.

### Mice

Young and aged C57BL/6 mice were obtained from the internal divisional stock (derived from mice obtained from The Jackson Laboratory). Young and aged acRFP C57BL/6 mice were obtained from the internal divisional stock (originally kindly donated by Prof. Fehling, Ulm University and previously described^89^). The NBSGW mice were obtained from the internal divisional stock (derived from mice obtained from The Jackson Laboratory, JAXStock No.026622) and were maintained as homozygotes. RhoAflox mice were described previously^20^ and crossed with CreErt2 mice (JAXStock No.008463). All mice were housed in the animal barrier facility under pathogen-free conditions at the Biomedical Research Institute of Bellvitge (IDIBELL). Throughout the manuscript, young mice are between 10 and 20 weeks old and aged mice are at least 80 weeks old. C57BL/6 mice were randomized for sex.

For the transplantation study NBSGW mice were randomized for sex and equal number of male and female mice were used across samples. Mice that failed to recover from blood sampling and mice that died due to laboratory errors were excluded. Mice that needed to be euthanized because they were scored as “weak and about to die” according to our approved animal license protocol for evaluating mouse health status remained part of the dataset. Allocation to control or treated group was done randomly (4-5 mice each experimental group in four different experimental batches). All animals were maintained according to the recommendations of the European Convention for the Protection of Vertebrate Animals used for Experimental and other Scientific Purposes (ETS 123). Animals were housed in groups of up to 4 animals per cage in Macrolon Type II (long) cages with bedding and paper nesting material. The animals had access to food (V1124-3, ssniff®) and water *ad libitum*. Animals were kept at a day/night rhythm of 12/12 hours throughout the entire experiment.

### Ethical compliance for mouse experiments

All mouse experiments were performed in compliance with the ethical regulations according to the Spanish Law for Animal Protection and Welfare Code and were previously approved in the project AR18008/10399 by IDIBELL’s Ethical Committee for Animal Experimentation (CEEA-IDIBELL) as well as by Generalitat of Catalunya.

### HSC transplantation in NBSGW

For HSC transplantation, young and aged acRFP C57BL/6 were used as donors. Two-hundred HSCs were sorted into 72-well Terasaki multi-well plates (Merck, M5812) and cultured for 16 h in HBSS + 10% FBS + 1% Penicylin/Streptomycin (P/S) with or without Rhosin (RhoA inhibitor or Ri)^43^ at 100µM in a water-jacketed incubator at 37 °C, 5% CO_2_, 3% O_2_. Cells were transplanted via retro-orbital vein injection. Peripheral blood chimerism was determined by FACS analysis every 6 weeks up to 18 weeks after transplants. The transplantation experiment was performed four times with a cohort of four recipient mice per group each transplant. In general, transplanted mice were regarded as engrafted when peripheral blood chimerism was higher or equal to 0.2% and contribution was detected in all lineages.

### Retroviral vector construction and viral production

H3K9 and the mutant form H3R9 sequences were kindly provided by the laboratory of Prof. Clemens Schmitt^97^. They are based on the H3.1 histone isoform with the K9 mutated to R9. Sequences were subcloned into the retroviral backbone pMSCVII (pMSCVII was a gift from Maki Nakayama (Addgene plasmid # 162750; http://n2t.net/addgene:162750 ; RRID:Addgene_162750) adding by PCR a P2A site and mCherry as a reporter. Resultant vectors were transfected into Phoenix-ECO using Lipofectamine 2000 (Thermo-Fisher) following manufacturer instructions. Supernatant containing retroviral particles was collected at 48h and 72h post-transfection and kept at 4°C until concentration 72h post-transfection. We used Millipore Centricon Plus-70, Ultracel-PL Membrane 100kDa (UFC710008 Millipore) for viral concentration. Concentrated retroviral particles were kept at -80°C prior to transduction. Retroviral particles were titrated in dilutions ranging from 1/10 to 1/500000 in NIH-3T3 cells (mouse embryonic fibroblasts). The titration was analyzed 48h later assessing mCherry-frequencies by flow cytometry (Cyto Flex SRT, MoFlo Cell Sorter). Viral infectious units (VIU) were calculated based on the initial cell-input, dilution, mCherry-frequency and volume of retroviral supernatant. The resulting values were plotted and the average, representing the transducing units (TU) was calculated from the linear portions of the graph.

### Flow cytometry and cell sorting

PB and BM cell immunostaining was performed according to standard procedures and samples were analysed by Beckman Coulter Gallios Analyzer (Beckman Coulter). RFP signal was used to distinguish donor from recipient cells. For PB and BM lineage analysis the antibodies used were all from eBioscience: anti-CD3ε (clone 145-2C11), anti-B220 (clone RA3-6B2), anti-Mac-1 (clone M1/70) and anti-Gr-1 (clone RC57BL/6-8C5). Lineage FACS analysis data are plotted as the percentage of B220^+^, CD3^+^ and myeloid (Mac-1^+^ and Gr-1^+^Mac-1^+^) cells among donor-derived RFP^+^ cells in case of a transplantation experiment or among total white blood cells within the bone marrow. As for early haematopoiesis analysis, mononuclear cells were isolated by low-density centrifugation (Histopaque 1083, Sigma) and stained with a cocktail of biotinylated lineage antibodies. Biotinylated antibodies used for lineage staining were all rat anti-mouse antibodies: anti-CD11b (clone M1/70), anti-B220 (clone RA3-6B2), anti-CD5 (clone 53-7.3) anti-Gr-1 (clone RB6-8C5), anti-Ter119 and anti-CD8a (clone 53-6.7) (all from eBioscience). After lineage depletion by magnetic separation (Dynabeads, Invitrogen), cells were stained with anti-Sca-1 (clone D7) (eBioscience), anti-c-Kit (clone 2B8) (eBioscience), anti-CD34 (clone RAM34) (eBioscience), anti-Flk-2 (clone A2F10) (eBioscience) and streptavidin (eBioscience). Early haematopoiesis FACS analysis data were plotted as percentage of long-term haematopoietic stem cells (HSCs, gated as LSK CD34^−/low^Flk2^−^), short-term haematopoietic stem cells (ST-HSCs, gated as LSK CD34^+^Flk2^−^) and lymphoid-primed multipotent progenitors (LMPPs, gated as LSK CD34^+^Flk2^+^)^98^ distributed among donor-derived LSKs (Lin^neg^c-Kit^+^Sca-1^+^ cells). To isolate HSCs, lineage depletion was performed to enrich for lineage-negative cells. Lineage-negative enriched cells were then stained as mentioned above and sorted using Beckman Coulter High Speed Cell Sorter Moflo-XDP (Beckman Coulter) and CytoFLEX SRT Cell Sorter (Beckman Coulter). For analysis of haematopoietic progenitors in the experiment of H3K9 and H3R9 transduction followed by transplantation, same procedure for BM analysis was carried out except for the staining after lineage magnetic depletion. Cells were stained with anti-IL7Ra (clone A7R34), anti-c-Kit (clone 2B8), anti-Sca1 (clone D7), anti-CD16/32 (clone 2.4G2) and anti CD34 (clone RAM34). Flow cytometry analysis data was plotted as percentage of MP (lin^-^, mCherry^+^, Il7Ra^-^, c-Kit^+^ and Sca1^-^), MEP (lin^-^, mCherry^+^, Il7Ra^-^, c-Kit^+^, Sca1^-^, CD16/32^-^, CD34^-^), CMP (lin^-^, mCherry^+^, Il7Ra^-^, c-Kit^+^, Sca1^-^, CD16/32^-^ CD34^+^), GMP (lin^-^, mCherry^+^, Il7Ra^-^, c-Kit^+^, Sca1^-^, CD16/32^+^ CD34^+^) and CLP (lin^-^, mCherry^+^, Il7Ra^+^, c-Kit^low^, Sca1^low^).

### LSK retroviral transduction and competitive transplantation in lethally irradiated mice

LSK (Lin^neg^c-Kit^+^Sca-1^+^) cells were sorted in growth medium (IMDM 10%FBS 1%P/S medium with cytokines mSCF, mTPO and G-CSF at 1µg/ml). They were seeded on fibronectin functionalized wells (50ng/ml) in 96 well plate, 10K cells per 50µl of medium and maintained in normoxia at 37°C and 5%CO_2_ for 20h. Then, medium was changed to growth medium containing polybrene (1/1000, Sigma-Aldrich) and 30MOI of the retroviral vector. Incubate the cells with the retroviral particles for 6-8h. Change medium to normal growth medium and incubate over night at 37°C, normoxia and 5%CO_2_. Cells were lifted with trypsin 0.05% for 5 minutes, quenched with medium with no cytokines, washed and recovered. A fraction of the cells was saved for transduction efficiency analysis and the rest was used for transplantation in lethally irradiated (9Gy) C57Bl6 mice. Cells for transplantation were mixed with BM competitor cells from non-irradiated C57Bl6 in a ratio 1/15 (20000 transduced LSK and 300000 competitor cells) in cold PBS.

### Immunofluorescence staining and confocal images acquisition

Freshly sorted HSCs were seeded on fibronectin-coated glass coverslips. For polarity staining, HSCs were incubated for 12–16h in HBSS + 10% FBS + 1% P/S and when indicated treated with 100 µM Rhosin^43^, 5mM NaB, 20 µM cPLA2 inhibitor (AACOCF3^23^), hypotonic medium or left untreated. After incubation at 37 °C, 5% CO_2_, 3% O_2_ in growth factor-free medium, cells were fixed with BD Cytofix fixation buffer (BD Biosciences). After fixation cells were gently washed with PBS, permeabilized with 0.2% Triton X-100 (Sigma) in PBS for 20 min and blocked with 10% donkey serum (Sigma) for 30 min. Primary and secondary antibody incubations were performed for 1 h at room temperature. Cells were stained with a DAPI dilution 1:500 in PBS of DAPI 1µg/µl (Thermo, ref) for 10 minutes at room temperature and washed twice with PBS. Coverslips were mounted with ProLong Gold Antifade reagent without DAPI (Invitrogen, Molecular Probes). A list of antibodies used for immunofluorescence staining is provided in Supplementary Data. Samples were imaged with an AxioObserver Z1 microscope (Zeiss) equipped with a ×63 PH objective. Alternatively, samples were analysed with an LSM880 confocal microscope (Zeiss) equipped with a ×63 objective. *Z*-stacks were obtained by automatically scanning along the *z* axis of the cell with a confocal microscope and acquiring a picture of the in-focus plane every 0.2-0.4 μm.

### Immunofluorescence Image Analysis and Rendering

Samples for immunofluorescence quantification analysis were rigorously sorted, stained and imaged in parallel within the same experiment to minimize any possible technical variability. Image acquisition at the confocal has been carried out consistently in between experiments regarding laser power, z-stack size and gain master. Antibody specificities have been validated using a knockout model in the case of RhoAGTP or using a control sample only with the secondary antibody in the staining protocol for the rest of the stainings. Quantification of protein signals has been done using Volocity Software 6.5 (Quorum technologies) using the “volume by intensity” tool, which sets a threshold for the positive signal against the negative. Positive signal threshold is set for each experiment by using RhoAKO for RhoAGTP and with the secondary antibody for the rest of the stainings. Morphometric measurements were done using Volocity Software 6.5 or by our computational vision approach. Quantifications of DAPI volume were done with the “volume” tool and Nuclear Height Average (NHA) and diameter was done using the tool “line”. Immunofluorescence images 3D reconstruction and rendering have been performed using Imaris 9.5.0 (Oxford Instruments) using the “surface” tool keeping the threshold constant for the signal in between experiments and conditions. As for polarity scoring, the localization of each single stained protein was considered polarized when a clear asymmetric distribution was visible by drawing a line across the middle of the cell. A total of 50 to 100 HSCs were singularly analysed per sample. Data are plotted as percentage of the total number of cells scored per sample.

### 3D nuclear DAPI images preprocessing

HSCs nucleus microscopy images were exported as Carl Zeiss CZI files for downstream analysis and processed using Python programming language. Images acquired with a different microscope to the ones specified above and images belonging to experiments in which more than 30 days passed between nucleus staining and image data acquisition were excluded from these analyses. As a first quality control, images displaying total pixel intensities higher than 4×108 were labeled as overexposed to DAPI and discarded. Similarly, images with increased Gaussian noise (estimated noise standard deviation > 8) were labeled as noisy and also excluded from further analyses. The remaining images were then corrected using Chambolle’s Total Variation denoising method^99^. Since image acquisition measurements were dynamically adjusted to the size of the nucleus, each 3D image was resized to achieve a uniform resolution of 10 pixels per micrometer in all dimensions through isotropic interpolation that accounted for variations in the number of z-stacks obtained. For each image, a nucleus binary mask was extracted using the Otsu segmentation method^100^, after smoothing with a Gaussian filter and allowing for hysteresis to preserve nuclear border with higher confidence Potential holes in the binary mask due to low intensity areas within the nucleus were filled and marked as part of the segmentation. Both the intensity image and respective nucleus mask were centered in the container array grid by trimming the background and padding the image borders symmetrically. Marginal intensities outside the nucleus mask were removed. To mitigate batch effects associated with technical variations in the fluorescence signal, we standardized the pixel intensity distribution within each nucleus mask using Z-score normalization to enhance comparability among conditions in downstream analysis.

### Intensity by distance to nuclear border analyses

Intensity by distance analyses were performed at iso-distance intervals of 0.1µm from the boundary of the nucleus segmentation, using a distance-transformed mask. A distance-transformed mask is a mask in which each pixel value represents the shortest geodesic distance to the nearest mask boundary. The average intensity value of all pixels within a 3D band with a thickness of 0.15µm was reported at each measurement interval. Young, aged and aged+Ri conditions were measured up to a distance of 1.6µm from the nuclear border. Boxplots of DAPI intensity by discrete distance ranges are computed taking into account the average intensity of all pixels within the specified distance boundaries for each interval.

### Multivariate feature analyses

Most morphometric and intensity features were measured with the scikit-image Python library^101^ regionprops_table() function at the nuclear mask level, DIR level, and on the largest 2D slide in the XY plane from each 3D image, comprising a total of 39 features **(Table S1).** The width, length, and height of each nucleus were obtained from its bounding box. Height deviation was calculated as the average standard deviation of height in the X dimension for each YZ slide. The aspect ratio was defined as the ratio of height to length. The surface area was calculated using the Marching Cubes algorithm after smoothing the nucleus mask with a Gaussian filter. The intensity ratio is computed as the ratio of average intensity within the 1 - 1.5 µm distance interval to the 0 - 0.5 µm distance interval from the nuclear border. The Excess of Perimeter (EOP) was computed as the proportion of the difference of the nuclear perimeter compared to the perimeter of an ellipse with the same major and minor axis length as the nucleus mask. Detailed information about each of these features is shown in **Table S1.**

DIRs were segmented using the Watershed method on the thresholded images. The intensity standardization of the images allowed for the choice of an absolute thresholding value for all nuclei. We set this parameter to the quantile 80% of the intensity distribution for all young nuclei. Individual DIRs were labelled and segmented with Watershed by identifying the intensity peaks as the local maxima in the Euclidean distance transform of the thresholded images. From the total of segmented DIRs, we filtered out those displaying a voxel volume smaller than 0.2µm. Morphometric and intensity features were measured on the resulting DIRs segmentation mask in the same manner as with the nuclear mask. DIRs distance to border was computed as the average of the distances for each voxel within each DIR to the nuclear mask border. After the calculation of these measurements, a second quality control is conducted to eliminate images that produced artifacts in the nuclear and DIR segmentation. The images were filtered out if they produced nuclear masks with a voxel volume smaller than 40µm^3^, an EOP larger than 0.25 or DIRs volume larger than 5µm3, which mostly belonged to confined nuclei that were damaged during the experimental setup and failed nuclear mask segmentations. We proceeded with 177 young nuclei, 164 aged nuclei, 144 aged+Ri nuclei.

Feature selection was performed on the original set of features to maximize the separation of samples according to our biological conditions of interest and minimize the information redundancy. First, statistically significant features which differ among young *vs.* aged and among aged *vs.* aged Ri-treated were found (Mann-Whitney U-test, p-value < 0.05). Later, features that exhibited absolute correlation values higher than 0.8 were discarded and the resulting sets from each pairwise comparison were merged to form a combined set of 20 features. These features were standardized and used to train the UMAP^52^ model on young, aged and aged Ri-treated nuclei. Clustering was conducted in the original multidimensional parameter space using the K-means algorithm^102^ with k=4 and depicted in the UMAP embedding. This value of k yielded the best silhouette scores for all clusters **(Figure S3F).** The resulting clusters revealed biologically relevant groups combining images from the young, aged, and aged+Ri conditions. Volcano plots showing feature importance for each of the identified clusters were generated by calculating the statistical significance (Mann-Whitney U-test) and log2 fold change of the average of each feature between the cluster and the average of the remaining clusters.

### Bone samples preparation for NanoIndenter

Samples were obtained from young and aged mice. Mice were perfused with PFA 4% in PBS, bones were collected as shown in^103^ and embedded in liquid OCT (CellPath, KMA-0100-00A) and frozen at -80°C. OCT is a cryo-embedding matrix, designed for cryostat sectioning at -10°C or below. For these experiments, humeri and femurs were used as they show an optimal ratio between rigidity of the bone walls and internal surface area. Samples for Chiaro NanoIndenter analysis were prepared by opening the bone structure and exposing the internal area of the bone marrow by gradually cutting transversally with a cryostat, keeping the sample at -20°C to avoid OCT thawing and exposing the bone marrow inner surface. The open bones were transferred to a 4-well Ibidi µ-slide (Ibidi, 80427) and partially embedded in agarose gel 4% (Condalab, 8010). The inner surface was covered with PBS to avoid tissue drying.

### Measurements with Chiaro NanoIndenter

For measuring stiffness of the tissue, Chiaro NanoIndenter was employed, and the data were elaborated through Piuma software (Optics11Life). 0.5 N/m - 25,6 µm-diameter probes (Optics11Life) were used for these experiments. The probes were calibrated through the Piuma software automated procedure, using a glass Petri dish filled with PBS. After the calibration, the Petri dish was replaced with the Ibidi µ-slide containing the samples of interest. The measurement was performed programming a 30-points matrix, a group of sequential measurements covering the whole diameter of the bone marrow. The points were distant in width (D_y_) 350 µm for the humerus and 450 µm for the femur; in length instead the distance (D_x_) was 150 µm for both humerus and femur. At least 3 matrixes per sample were performed, two on the peri-epiphyseal part and one on the diaphyseal part. The effective Young’s modulus was derived from force vs. indentation curves, using a Hertzian model. The stiffness of the bone marrow was measured and calculated in kPa (kiloPascals) units.

### Protein Sample preparation

Bone marrow cells for western blot were isolated by flushing bones from young and aged mice into HBSS (Gibco, 24020117), complemented with FBS 10% and P/S 1%. Cells were then centrifuged at 1500 RPM for 5 minutes at RT, resuspended in 3 ml HBSS per mouse and filtered through a 70 µm cell strainer. For each mouse, 27 ml of 1X Red Blood Lysis Buffer (Biolegend, 420301) were added and incubated for 5 minutes at RT in the dark. At the end of the incubation, PBS was added to the top of the tube, to stop the lysis reaction and dilute the buffer. Cells were then centrifuged at 1500 RPM for 5 minutes at 4°C, washed twice and resuspended in PBS. After cell count, cells were pelleted, washed with PBS and lysed with RIPA buffer (Pierce™ RIPA Buffer, ThermoFisher, 89900) complemented with protease inhibitor cocktail (cOmplete, Sigma Aldrich, 11697498001), for 15 minutes on ice, flicking the tube several times during the incubation. The cells were then centrifuged at 14,000 RPM for 15 minutes at 4°C; the supernatant containing the protein lysate was stored at -80°C.

### RhoA-GTP bound Pull Down

To isolate the active isoform of RhoA, GTP-bound RhoA, Active RhoA Detection Kit (Cell Signaling, 8820) was used following manufacturer instructions.

### Western Blotting

Acrylamide gel was prepared, run and transferred in nitrocellulose membranes following the indications of the kit manufacturer (TGX™ FastCast™ Acrylamide Kit 12%, BioRad, 1610185). The transferred membrane was cut as needed and washed twice in TBS-T (Tris-buffered saline (TBS) 1x Tween20 0.1%) for 15 minutes each time, then blocked with low-fat milk 5% in TBS-T on a rocker for 1 hour at RT. Anti-RhoA (rabbit monoclonal, Cell Signaling, 2117) and anti-actin (mouse monoclonal, Sigma Aldrich, A2228-100UL) primary antibodies were diluted 1:100 and 1:2000 respectively in milk 5% in TBS-T and incubated in a cold room with the samples at 4°C overnight. After incubation, the membranes were washed twice in TBS-T for 15 minutes each, and incubated with HPR-goat anti-rabbit (Biorad, 1706515) and anti-mouse (Biorad, secondary antibodies (BioRad), diluted in milk 5% in TBS-T, at a concentration of 1:2000 and 1:5000 respectively, for 1 hour at RT. The membranes were then washed twice in TBS-T for 15 minutes. ECL Prime Western Blotting Detection Reagents (GE Healthcare, 28980926) was used for developing, by pipetting 500 µl of reagent A and B on the membrane and incubating it for 5 minutes in the dark. The antibodies were detected by UV reveal in BioRad’s ChemiDoc and analyzed using Image Lab 6.1 software (BioRad).

### Confinement assay

The confinement device used in this protocol was adapted to a single well plate from a previously described method^25,104^. Briefly, the confinement device was made by a magnetic container, two metallic rings, a polymeric ring and a closing ring (**Figure S1A**). The compression is mediated by a pillar coverslip, a polymeric piston and the magnetic lid of the device that exerts the confinement pressure. 3,000 HSCs were seeded in a volume of 40 µl HBSS 10% FBS 1% P/S onto a 35-mm glass coverslip. The coverslip was previously functionalized with fibronectin: 40 µl of fibronectin (50 µg/ml) were applied upon the surface of the coverslip for 2 hours at RT, blocked with the same volume of BSA 2% for 1h at 37°C and washed with PBS Ca^2+^/Mg^2+^. The coverslip with cells was mounted in the confinement device and compressed with pillars of 3, 5 and 8 µm for 2 hours at 37°C in a hypoxic incubator (5% CO_2_, 3% O_2_). Cells were then fixed directly in the confinement device with 1 ml of PFA 4.21% (BD Cytofix, Thermo Fisher, 15817828) at 4°C for 15’. The sample on the coverslip was then removed from the confiner and stocked at 4°C in PBS for a maximum of 2 weeks before staining.

### Hypotonic shock assay

Briefly, HSCs were cultured on fibronectin functionalized coverslips in 30 µl of isotonic medium (HBSS 10% FBS 1% P/S) for 12h after sorting in a hypoxic incubator (5% CO_2_, 3% O_2_). Then, the medium was removed almost completely carefully and replaced by fresh isotonic medium or by hypotonic medium. Hypotonic medium consists on a 0.75X dilution of isotonic medium with sterile distillated water. HSCs were incubated for 2h in the same hypoxic conditions and then fixed as explained above. Protocol of staining was performed as usual.

### Nuclear Wrinkling Analysis

To analyze nuclear wrinkling in LaminB stained HSCs, we developed an automatic image analysis pipeline in Matlab (Mathworks, R2024b). Fluorescence images of LaminB were acquired in a LSM880 confocal microscope (Zeiss) commented in section above “*Immunofluorescence staining and confocal images acquisition”* For each dataset, the equatorial maximum cross-section section was identified from Z-slices and 3 slices were selected above and below, covering approximately 50 percent of the nucleus surface with optimal resolution of surface wrinkles in the lateral X-Y dimension. Selected image slices were then subjected to binning (factor n=3) using a bilinear interpolation to reduce noise and facilitate subsequent processing steps, effectively preserving key structural information. A threshold intensity of 25 counts was applied to create a binary mask and to segment the regions of interest representing the nuclear envelope. A Gaussian blur with a sigma of 3 pixels was applied to the binary mask to smooth the segmented regions and facilitate the subsequent skeletonization of nucleus envelope features based on a morphological thinning algorithm. The skeletonization process enabled to extract the structural features of the segmented Lamin signal, highlighting the wrinkles of the nuclear envelope. To remove structural artefacts outside the nuclear region of interest, the skeletonized image was filled to generate an inner region (representing the inside of the nuclear wrinkles) and an outer region (around the nucleus periphery) by applying an inverted mask to the filled skeleton. This allowed to separate the nuclear envelope from the surrounding structures and detect the points representing nuclear invaginations. The coordinates of the nuclear surface and inner invaginations represent the total nuclear periphery and were extracted for further quantitative metrics of nuclear wrinkles. We derived the nuclear circularity (𝐶 = 4𝜋𝐴⁄𝑃^2^) from the total perimeter 𝑃 and area 𝐴 of nuclear cross-sections from multiple Z-slices for each cell. The circularity provides a quantitative measure of nuclear envelope deformations, with values close to 1 indicating a more circular shape and values < 1 a highly deformed and wrinkled nuclear surface. In addition, we derived the excess folding parameter (𝐸 = 1 − 𝑝⁄𝑃) as the ratio of the boundary outline of the nucleus 𝑝 to the total perimeter 𝑃 of nuclear cross-sections, with values close to 0 resembling a circular shape and values closer to 1 indicating increased nuclear wrinkling. Statistical data analysis was performed in Prism (Version 10.2.3.) using a Kruskal-Wallis test for multiple comparisons.

### Bulk ATAC-seq of HSCs

HSCs were isolated from young and aged animals via FACS sorting. Between 2,000 and 5,000 cells were isolated for each library and we prepared 3-5 biological replicates for each sample arm. Cells were cultured overnight without growth factors at 3% O2 and washed twice with PBS before processing. Aged cells from same animal were used as aged control sample and aged treated with Ri sample. Young cells from same animal were used as young control sample and young confined under 5μm sample. Cells were subjected to fragmentation of open chromatin regions using Tn5 transposase (Illumina), followed by a pre-amplification step, library preparation and subsequent paired-end sequencing. For the pre-amplification, NEBNext Ultra II Q5 Master Mix was used with Primer 1: 5’GTCTCGTGGGCTCGGAGATGTGTATAAGAGACAG3’ and Primer 2: 5’TCGTCGGCAGCGTCAGATGTGTATAAGAGACAG3’. For dual-indexing, 10 μL of the pre-amplified ATAC reaction was used as input for Nextera XT index kit (Illumina). The generated libraries were quantified using an Agilent Bioanalyzer and a qPCR kit (New England Biolabs), pooled and subjected to next-generation sequencing in a NextSeq550 or Illumina HiSeq 2000 for paired-end 150 bp or 250 bp sequencing condition. Initial quality control was performed with FastQC v0.11.5. The ENCODE ATAC-seq pipeline v2.1.3 was used to process the FASTQ files, including adapter trimming, alignment to the reference genome (mm10), filtering and peak calling, and using the parameters “atac.pipeline_type” : “atac”, “atac.align_only” : false, “atac.true_rep_only” : false, “atac.paired_end” : true, “atac.auto_detect_adapter” : true, “atac.multimapping” : 20, “atac.mapq_thresh” : 20. Samtools v1.14^105^ was used to index BAM files. ATACseqQC v1.20.2^106^, BSgenome v1.64.0, Bsgenome.Mmusculus.UCSC.mm10 v1.4.3 and TxDb.Mmusculus.UCSC.mm10.knownGene v3.10.0 in R v4.2.0 and Bioconductor v3.15.2 were used for quality control and Tn5 shifting. Deeptools v3.5.1^107^ was used to transform BAM files to normalized BigWig files (with parameters --effectiveGenomeSize 2407883318 --normalizeUsing RPKM --exactScaling --binSize 50 –extendReads) to be visualized in the Integrative Genomics Viewer (IGV)^108^ web app and to check for read enrichment in transcription start sites (TSS). A consensus set of peaks was defined by taking those peaks detected in at least 6 samples. This threshold was determined using Monte Carlo simulation. Briefly, a binary matrix was generated including all detected peaks as rows and all samples as columns, with 1 for presence of the peaks and 0 for absence. The binary matrix was randomized 1,000 times and the number of peaks detected in all possible minimum number of samples was calculated for each randomized matrix. The chosen minimum number of samples was the one where the mean number of consensus peaks in the simulated data was ≈10% of the number of consensus peaks in the empirical data (false positive rate, FPR). The final 57,289 consensus peaks were resized to have a width of 700 bp. In the case of the young 5μm confined analysis, 42,632 consensus peaks were defined following the same strategy but using only the young and aged control samples. Peaks were annotated using annotatePeaks function from Homer v4.11.1^109^ and a donut chart was plotted using ggplot2 v3.3.6, RColorBrewer v1.1-3 and ggrepel v0.9.1.

In R v4.2.0 and Bioconductor v3.15.2, the featureCounts function from Rsubread v2.10.5^110^ was used to count the number of reads in consensus peaks with parameters largestOverlap = TRUE, isPairedEnd = TRUE, countReadPairs = TRUE, requireBothEndsMapped = TRUE, checkFragLength = TRUE, minFragLength = 0, maxFragLength = 2000. The count matrix was transformed and normalized using the voom function from limma v3.52.1^111^ and the quantile normalization method. Batch effects were removed with the removeBatchEffect function only for visualization purposes. Principal component analysis (PCA) was performed using the PCA function from FactoMineR v2.6^112^ and the top 10,000 most variable peaks. limma was used to perform differential accessibility (DA) analysis between conditions, adding the batch as a covariate. The p-value threshold to determine the significance of the DA was defined using Monte Carlo simulation. Briefly, the normalized count matrix was randomized 1,000 times and the number of DARs at different p-value thresholds was calculated for each randomized matrix. The chosen p-value threshold was the one where the number of DARs in the simulated data was ≈6-8% of the number of DARs in the empirical data (FPR), a p-value of 0.005 in the case of the Ri analysis and 0.001 in the case of the young 5μm confined analysis. Volcano plots were plotted using ggplot2, ggrepel and patchwork v1.1.1. Venn diagram was plotted using VennDiagram v1.7.3^113^. Heatmap was plotted using pheatmap v1.0.12. GO enrichment analysis of genes close to DARs (considered close if the distance of the DAR to the TSS of the gene was less than 100 kb upstream or 25 kb downstream) was performed using clusterProfiler v4.4.1^114^, AnnotationDbi v1.58.0 and org.Mm.eg.db v3.15.0 with the function enrichGO and parameters ont = “BP”, pvalueCutoff = 0.1, pAdjustMethod = “BH”, minGSSize = 10, maxGSSize = 500, readable = TRUE. Selected GOs were plotted in radar plots using the function radarchart from fmsb v0.7.4. Barplots representing log2FCs in the proportions of genomic region types among the DARs from the different comparisons compared to the proportions in the consensus peaks were plotted using ggplot2, RColorBrewer and patchwork. Fisher’s exact tests were performed to compare the proportions in DARs vs the proportions in consensus peaks. One-proportion z-tests were used to compare the proportion of a specific genomic region (i.e. intron) among the DARs vs among the consensus peaks. TF-binding motifs in the different sets of DARs were found using the findMotifsGenome function from Homer, using the consensus peaks as background. In R v4.3.0 and Bioconductor v3.17, ATACseqTFEA v1.2.0 was used to obtain the coordinates of the Klf4 motif (Jaspar MA0039.4) across the DARs opening with Ri, which were later used to determine the Klf4-targeted genes (<100kb upstream or <25kb downstream).

### Bulk RNA-seq of HSCs

RNA-seq libraries were generated from 2,000 pooled young and aged HSCs (n=3 biological repeats per condition). Cells were cultured overnight without growth factors at 3% O2 and washed twice with PBS before processing. Aged cells from same animal were used as control sample and aged treated with Ri sample. SMART-Seq® v4 Ultra® Low Input RNA Kit for Sequencing manufacturer’s protocol was strictly followed. We quantified the cDNA quality and quantity using an Agilent Bioanalyzer. For the library preparation, 150 pg of cDNA were used per sample using the NexteraXT index kit. We performed quality control of our libraries using an Agilent Bioanalyzer and quantifying with qPCR kit (New England Biolabs). Libraries were then pooled to be subjected to next generation sequencing in a NextSeq550 for paired-end 150 bp sequencing condition.

After performing quality control with FastQC v0.11.5, adapters were removed from the FASTQ files using Cutadapt v1.18^115^ with parameters -m 20 -O 6 -q 20. Reads were mapped to the reference genome mm10 (Mus Musculus GRCm38, Ensembl 102 Nov 2020) using STAR v2.7.0^116^ and BAM files were sorted and indexed using Samtools v1.14^105^. Library complexity was estimated using Picard v2.26.7. Deeptools v3.5.1^107^ was used to transform BAM files to normalized BigWig files (with parameters -- effectiveGenomeSize 2407883318 --normalizeUsing CPM --exactScaling --binSize 50) to be visualized in IGV^108^ web app and to check for read enrichment in exons.

In R v4.2.0 and Bioconductor v3.15.2, the featureCounts function from Rsubread v2.10.5^110^ was used to count the number of reads in genes for each sample, using Ensembl GTF annotation for GRCm38 102 version, filtered for protein coding genes and with parameters GTF.featureType = “exon”, GTF.attrType = “gene_name”, useMetaFeatures = TRUE, isPairedEnd = TRUE, countReadPairs = TRUE, requireBothEndsMapped = TRUE, countMultiMappingReads = TRUE, fraction = TRUE. The function filterByExpr from edgeR v3.38.1^117^ was used to keep only those genes with more than 20 counts in at least three samples and a minimum total count of 60. The count matrix was transformed and normalized using the voom function from limma v3.52.1^111^ and the quantile normalization method.^112^ limma was used to perform differential expression (DE) analysis between the sequencing batches and 24 genes were removed from the analysis for showing significant batch effect (with <5% FPR; p-value < 0.0001). The DE analysis between conditions was also performed with limma. The p-value threshold to determine the significance of the DE was defined using Monte Carlo simulation as for the DA analysis in ATAC-seq. In this case, the chosen p-value threshold was the one where the FPR was ≈5%, a p-value of 0.001. Volcano plots were plotted using ggplot2 v3.3.6, ggrepel v0.9.1 and patchwork v1.1.1. Venn diagram was plotted using VennDiagram v1.7.3^113^. GSEA was performed using clusterProfiler v4.4.1^114^, AnnotationDbi v1.58.0 and org.Mm.eg.db v3.15.0 with the function gseGO and parameters ont = “BP”, pvalueCutoff = 0.05, pAdjustMethod = “BH”, minGSSize = 10, maxGSSize = 500, seed = TRUE, eps = 0. Selected GOs were plotted in radar plots using the function radarchart from fmsb v0.7.4 and the enrichment curve for “acute inflammatory response” was plotted using the gseaplot2 function of enrichplot v1.16.1. GSEA for the Interferome.org^71^ (filtering by Species: Mus musculus; System: Haemopoietic/Immune; Cell: HSC or haematopoietic stem cells; and FC: 2), interferon-stimulated^72^ and aging^73^ signatures was performed with the function GSEA of clusterProfiler and parameters minGSSize = 1, maxGSSize = Inf, pvalueCutoff = Inf, pAdjustMethod = “BH”, seed = TRUE and plotted using the gseaplot2 function of enrichplot.

To analyse REs, the STAR alignment was repeated with parameters --winAnchorMultimapNmax 200 -- outFilterMultimapNmax 100 to allow for more multimapping. The function TEcount from TEtranscripts v2.2.1^118^ was used to count the reads mapping to RE subfamilies, using the annotation downloaded from Hammell’s lab website for GRCm38 Ensembl rmsk. The resulting count matrix was processed as previously described for the genes but keeping only those RE subfamilies with more than 10 counts in at least three samples and a minimum total count of 30. No RE subfamilies were significantly affected by the sequencing batch. The DE significance p-value threshold used was 0.005 (FPR ∼8%). Venn diagram was plotted using VennDiagram. Heatmap was plotted using ComplexHeatmap v2.12.1^119^. Boxplot was plotted using ggplot2.

The t-statistics for the Klf4-targeted genes were plotted with ggplot2 and ggrepel. GO enrichment of upregulated Klf4-targeted genes was performed using clusterProfiler, AnnotationDbi and org.Mm.eg.db with the function enrichGO and parameters ont = “BP”, pvalueCutoff = 0.05, pAdjustMethod = “BH”, minGSSize = 10, maxGSSize = 500, readable = TRUE. Selected GOs were plotted in a radar plot using the function radarchart from fmsb.

### Single-cell RNA seq

LSK cells were sorted as explained previously from young and aged mice (n = 3 for each). Cells were incubated 16h in IMDM 10%FBS 1%P/S in the presence or absence of Ri 100µM and then profiled by using standard 10x Genomics protocols for single cell sequencing. After performing the quality control of the FASTQ files with FastQC v0.11.5, reads were aligned to the reference genome (GRCm38/mm10, annotation Ensembl 98), filtered, and counted using Cell Ranger software v7.2.0 (10x Genomics). Count matrices were pre-processed using R v4.2.0 and Seurat package v4.1.1^120^. In an initial filtering step, genes expressed in <10 cells and cells expressing <10 genes were discarded. Cells with >6% of mitochondrial RNA, <1,500 genes, >7,000 genes, <1,500 UMI counts, or >40,000 UMI counts were also discarded. We further discarded genes with less than 400 UMI counts (decided based on the distribution of the total counts per gene). We obtained a total of 60,648 cells and 15,049 genes to continue with the analysis. After log-normalizing the data, the genes defined by Kowalczyk et al.^121^ associated to G0, early G1, late G1, S and G2/M cell cycle phases were used to score each cell for the average expression of each set of genes using the AddModuleScore function of Seurat. A cell cycle phase was assigned to each cell according to the highest score. Seurat’s SCTransform function was used to normalize the gene counts for each condition and sequencing batch separately, regressing out the effect of the number of genes, the number of UMI counts, the percentage of mitochondrial RNA, and the scores for the cell cycle phases in every cell and returning 3000 variable genes for each condition. Then, integration was performed following Seurat’s integration workflow, using 3000 integration features and canonical correlation analysis with 30 dimensions. The integrated dataset, 50 principal components (PCs), and 500 epochs were used to generate the UMAP. Clustering was performed with Seurat’s FindNeighbors and FindClusters functions, using 50 PCs, Louvain algorithm, and a resolution of 0.2, chosen after evaluating several resolutions with the clustree package v0.4.4^122^. The resulting clusters were annotated based on their markers (obtained after running the FindAllMarkers function on the log-normalized data with parameters test.use = ‘wilcox’, logfc.threshold = 0.25, min.pct = 0.1, only.pos = TRUE, return.thresh = 0.05), plotting the expression of genes known to be expressed in HSPCs, and projecting on our data different gene signatures defined in previous studies (like the MolO signature defined by Wilson et al.^82^, the low/high-output HSC signature defined by Rodriguez-Fraticelli et al.^83^, and the dormant/active HSC signature defined by Cabezas-Wallscheid et al.^84^) using the AddModuleScore function of Seurat. Plots were generated using Seurat’s plotting functions and ggplot2 v3.3.6. scCODA^123^ v0.1.9 was used to perform compositional analysis, using the cluster CyclingCells_2 as the reference (as automatically selected by the tool), and running the model 10 times to account for the randomness introduced by the MCMC sampling. Differences were considered statistically credible if at least 7 of the runs showed an effect. In the HSC cluster, differential expression analysis between the three conditions was performed using muscat^124^ v1.10.1 (Bioconductor v3.15.2) and the muscat_analysis function of the muscatWrapper v1.0.0 R package on the log-normalized data with parameter de_method_oi = “limma-voom” and regressing out the effect of the sequencing batch. Significance was considered if |log2FC| > 1 and FDR < 0.05. MAplots were plotted using ggplot2 and ggrepel v0.9.1. To perform the GSEA of the signature for hemogenic precursors defined by Pereira et al.^85^, we first ordered the genes by their -log10(p-value) * sign(log2FC). Then the GSEA function from clusterProfiler v4.4.1^114^ was used to determine the enrichment of the signature with the parameters minGSSize = 1, maxGSSize = Inf, pvalueCutoff = Inf, pAdjustMethod = “BH”, seed = TRUE, eps = 0. The p-values were adjusted across the three pairwise comparisons between the conditions. The enrichment plot was generated using the gseaplot2 function of the package enrichplot v1.16.1. SCENIC v1.3.1^125^ was used to calculate regulons (TF + target genes) activity per cell in the HSC cluster. These 225 regulon activity scores were used to integrate the data by condition and sequencing batch following Seurat’s integration workflow, using all regulons as integration features and canonical correlation analysis with 30 dimensions. The integrated data was scaled and centered using the ScaleData function of Seurat and the effect of the number of genes, the number of UMI counts, the percentage of mitochondrial RNA, and the scores for the cell cycle phases in every cell was regressed out. The UMAP was generated using 25 PCs and 500 epochs. The activity scores of interesting regulons were binarized based on the distribution of their values (AUC = 0.128, 0.007 and 0.01 for Klf4, Ctcf and Hoxa10, respectively). Differential regulon activity analysis between conditions was performed using muscat^124^ and the muscat_analysis function of the muscatWrapper R package on the AUC values with parameter de_method_oi = “limma-voom” and regressing out the effect of the sequencing batch. Significance was considered if |log2FC| > 0.5 and FDR < 0.05. The activity scores of the significantly different regulons were plotted in a heatmap using ComplexHeatmap v2.12.1^119^.

## Notes

### Competing Interest Statement

The findings presented in this study are covered under patent application number EP25382180.5, filled on 27/02/2025.

