## Supplementary figures and images for "Targeting RhoA activity rejuvenates aged hematopoietic stem cells"

### Supplementary Figures

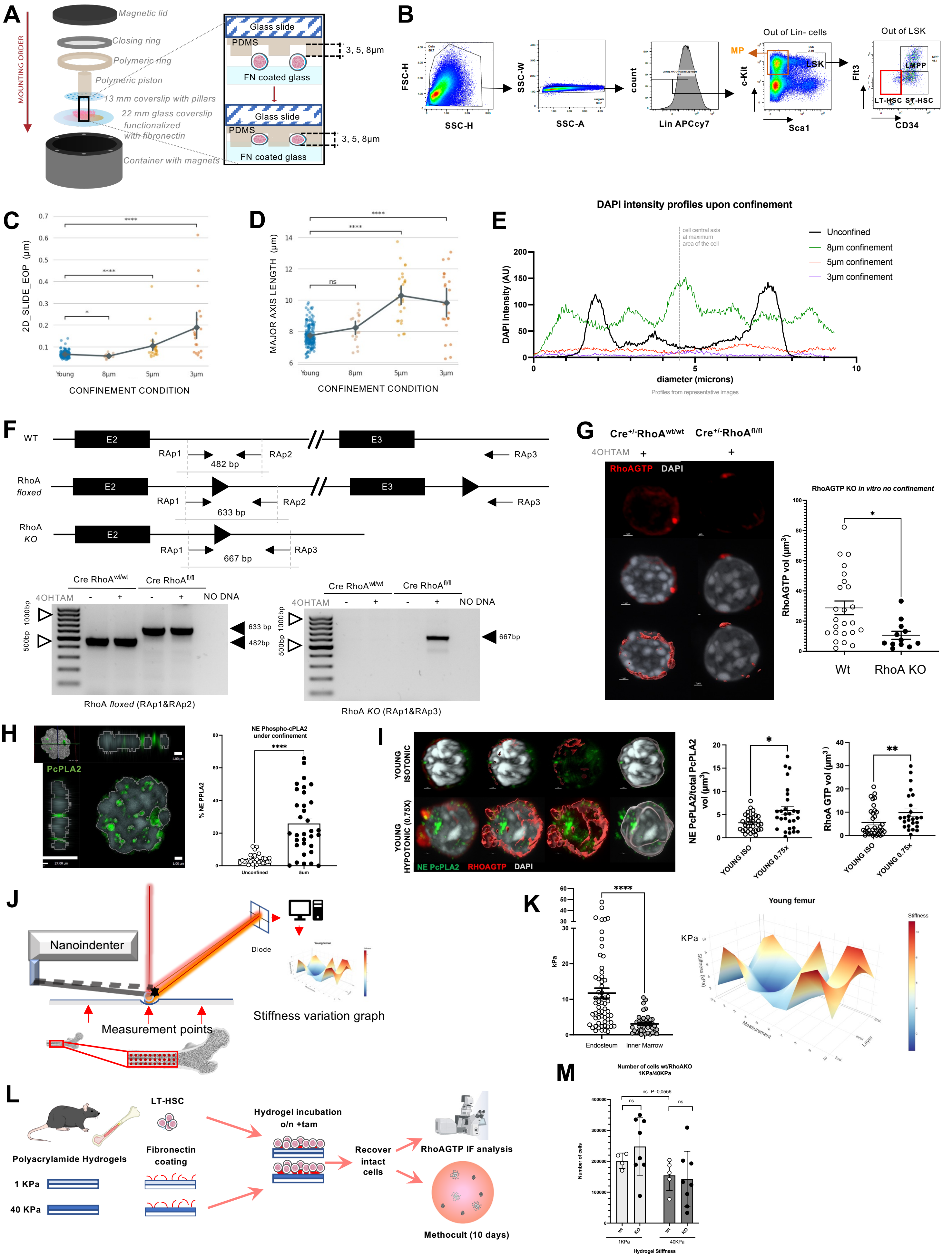

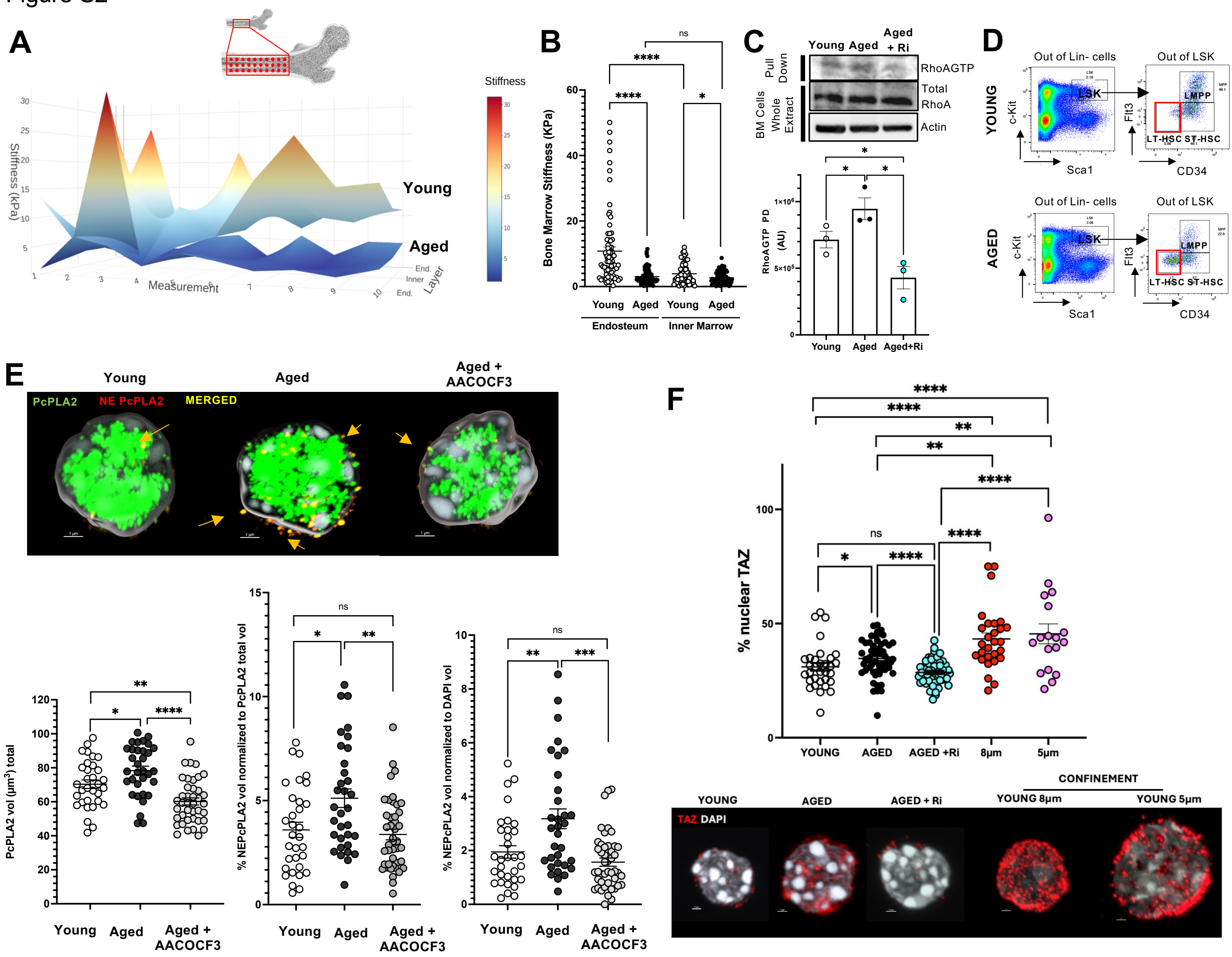

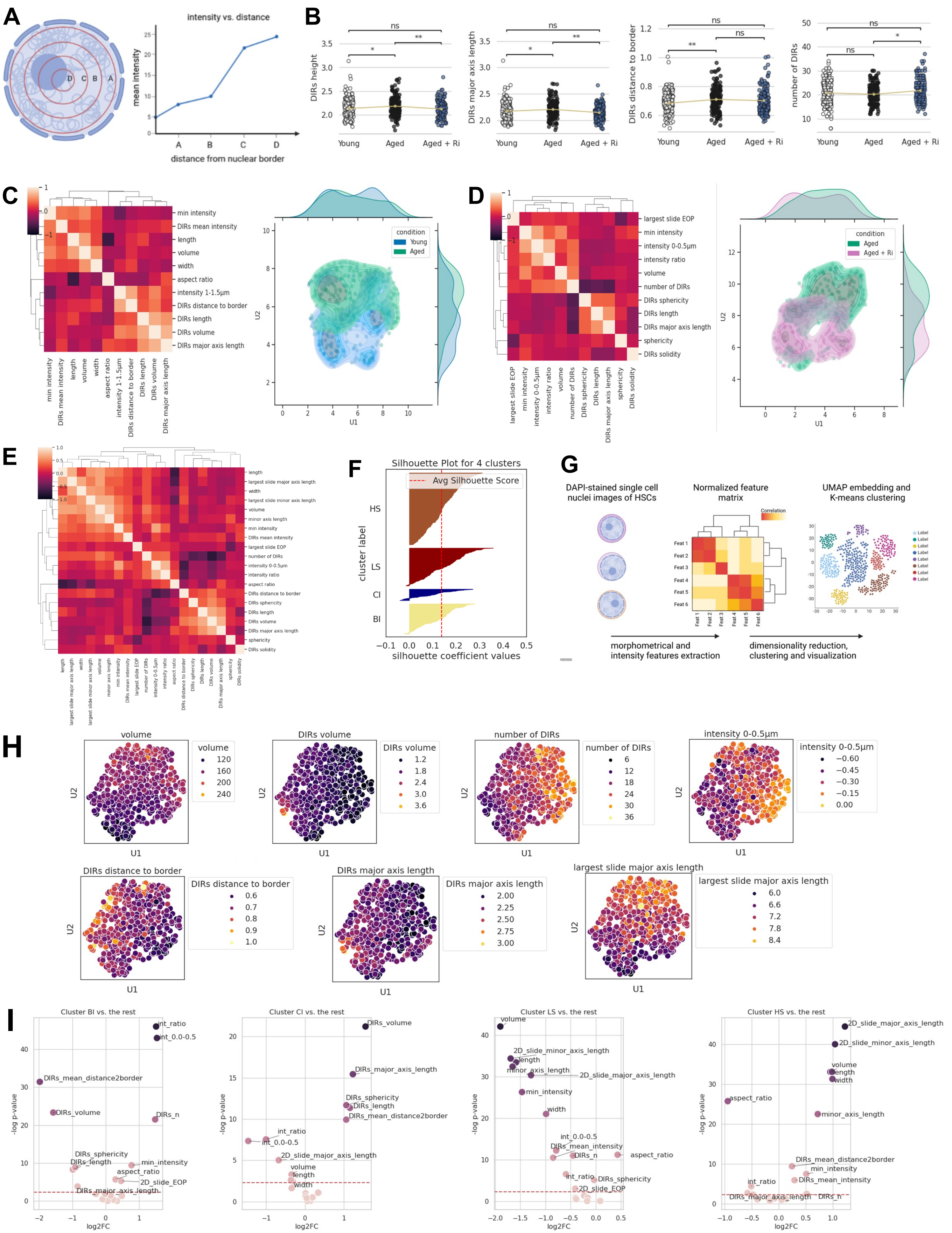

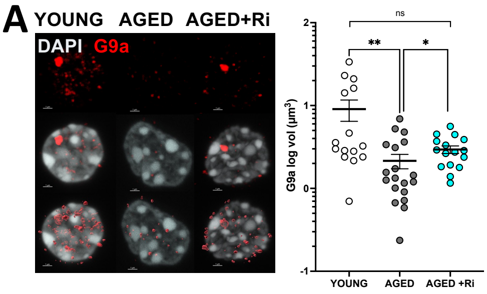

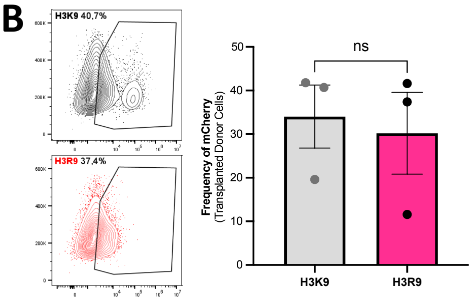

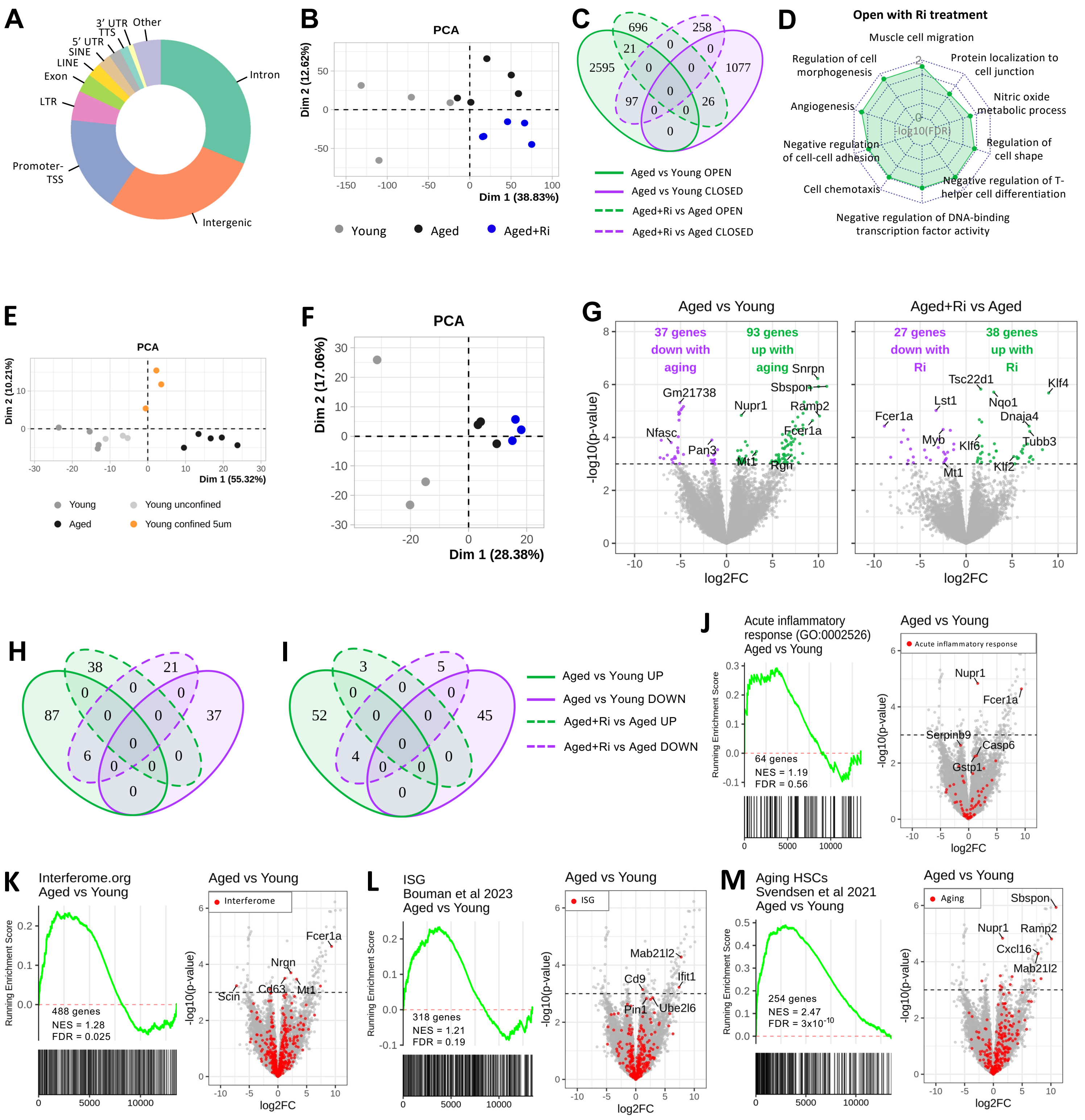

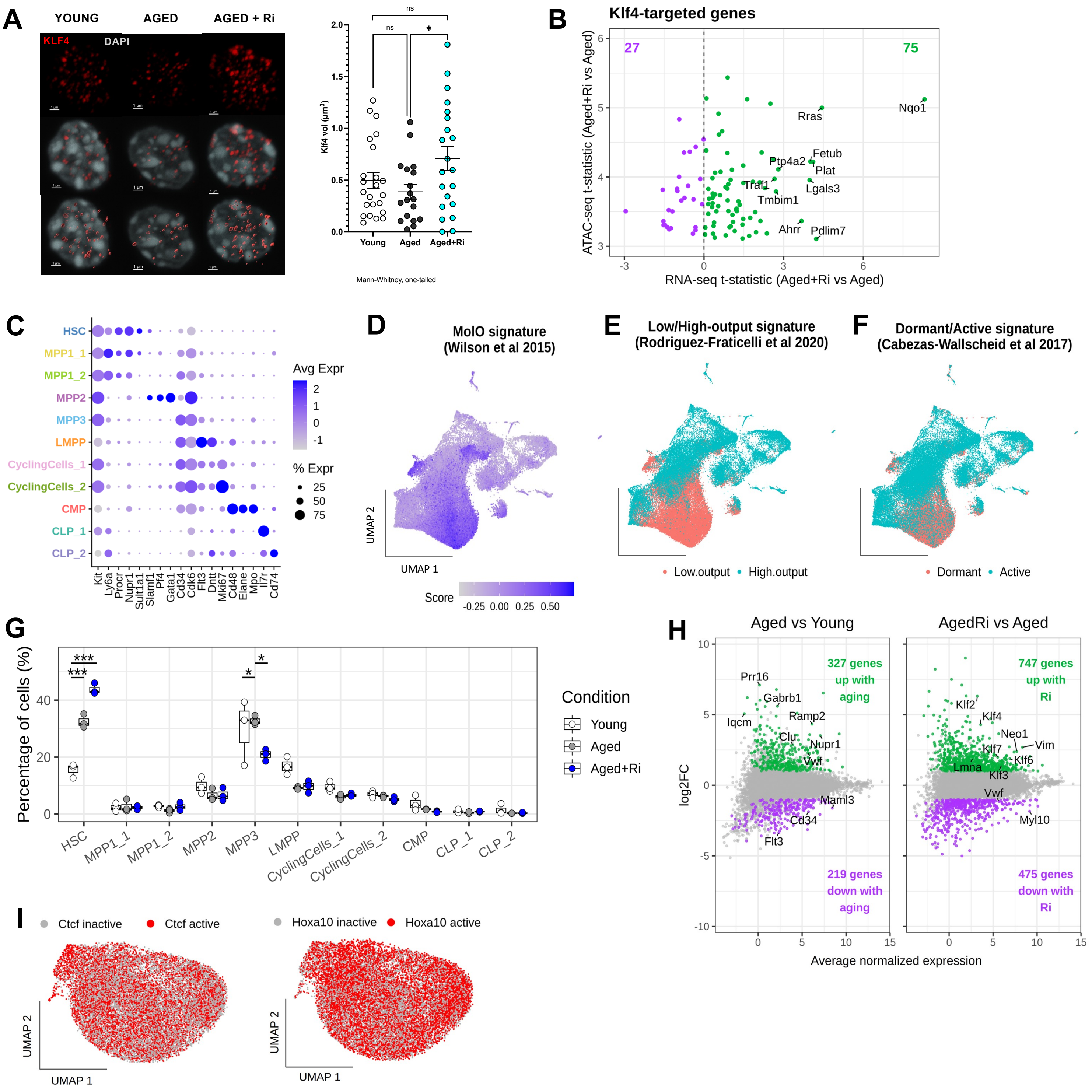

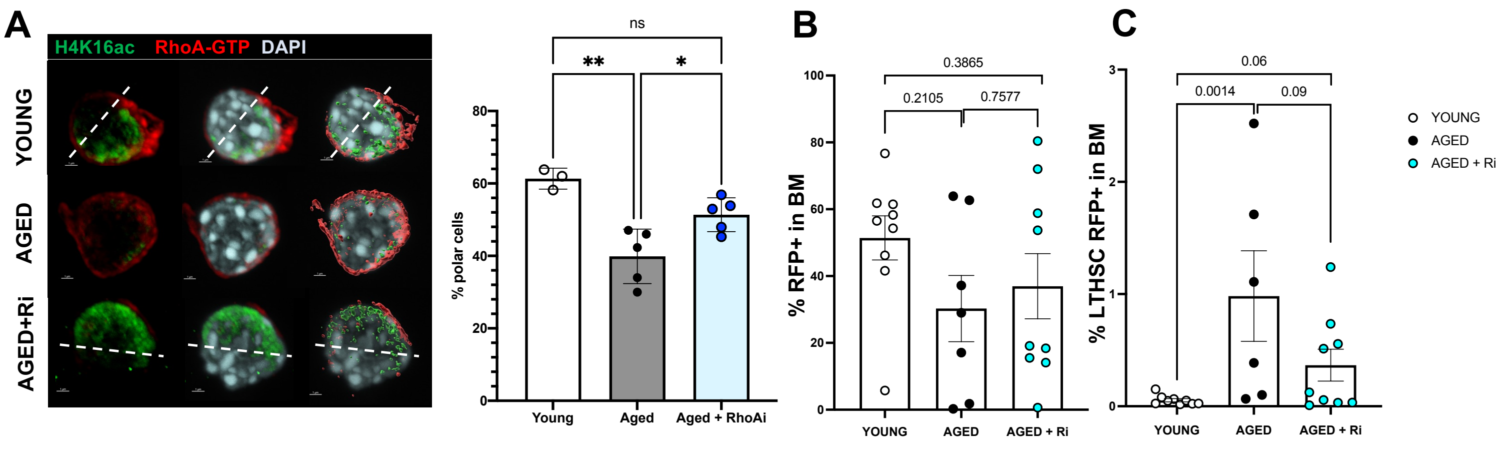
