## Supplementary material for "Targeting RhoA activity rejuvenates aged hematopoietic stem cells": Table S1

**Supp. Table 1**

| Variable ID | Variable Name | Units | Description |
|---|---|---|---|
| volume | Nuclear Volume | $\mu m^3$ | The number of pixels within the nucleus mask in 3D |
| width | Nuclear Width | μm | The maximum size of the nucleus mask in the X dimension, obtained from the nucleus mask bounding box |
| length | Nuclear Length | μm | The maximum size of the nucleus mask in the Y dimension, obtained from the nucleus mask bounding box |
| height | Nuclear Height | μm | The maximum size of the nucleus mask in the Z dimension, obtained from the nucleus mask bounding box |
| height deviation | Nuclear Height Deviation | μm | Mean standard deviation of the height measurements for each YZ slide in the 3D nucleus mask |
| aspect ratio | Aspect Ratio | - | The ratio between the height and the length of the nucleus mask |
| major axis length | Length of Major Axis | μm | The length of the major axis of the ellipse that has the same normalized second central moments as the nucleus mask |
| minor axis length | Length of Minor Axis | μm | The length of the minor axis of the ellipse that has the same normalized second central moments as the nucleus mask |
| equivalent diameter | Nuclear Equivalent Diameter | μm | The diameter of a sphere with the same volume as the nuclear volume |
| max intensity | Maximum Intensity | - | The value of the highest intensity in the nucleus mask |
| min intensity | Minimum Intensity | - | The value of the lowest intensity in the nucleus mask |
| sphericity | Sphericity | - | The roundness of the nucleus mask relative to a sphere in 3D. Computed as indicated in equation 1 below. |
| surface area | Nuclear Surface Area | - | The ratio of the number of pixels in the nucleus mask to the number of pixels in the nucleus bounding box |
| DIRs volume | DAPI-Intense Regions Volume | $\mu m^3$ | The mean of the number of pixels within each DIRs in the DIRs mask |

| | | | |
|---|---|---|---|
| DIRs width | DAPI-Intense Regions Width | µm | The average for all DIRs of the maximum size of the DIR mask in the X dimension, obtained from the DIR mask bounding box |
| DIRs length | DAPI-Intense Regions Length | µm | The average for all DIRs of the maximum size of the DIR mask in the Y dimension, obtained from the DIR mask bounding box |
| DIRs height | DAPI-Intense Regions Height | µm | The average for all DIRs of the maximum size of the DIR mask in the Z dimension, obtained from the DIR mask bounding box |
| DIRs aspect ratio | DAPI-Intense Regions Aspect Ratio | - | The mean of the ratios between the height and the length of all the DIRs within the DIRs mask |
| DIRs surface area | Area of the Surface of the DAPI-Intense Regions | $\mu m^2$ | The mean of the area of the surface of all the DIRs within the DIRs mask. The surface area was calculated using the Marching Cubes algorithm after smoothing the nucleus mask with a Gaussian filter. |
| DIRs major axis length | Length of Major Axis of the DAPI-Intense Regions | µm | The mean from all DIRs of the length of the major axis of the ellipse that has the same normalized second central moments as each DIR in the DIRs mask |
| DIRs minor axis length | Length of Minor Axis of the DAPI-Intense Regions | µm | The mean from all DIRs of the length of the minor axis of the ellipse that has the same normalized second central moments as each DIR in the DIRs mask |
| DIRs sphericity | Sphericity of the DAPI-Intense Regions | - | The mean of the roundness of each DIR in the DIRs mask relative to a sphere in 3D. Computed as indicated in equation 1 below |
| DIRs mean intensity | Mean Intensity of the DAPI-Intense Regions | - | The mean of the average intensity value for each DIR in the DIRs mask |
| DIRs solidity | Solidity of the DAPI-Intense Regions | - | The mean of the ratios of pixels in the region to pixels of the convex hull image of each DIR in the DIRs mask |
| DIRs distance to border | Mean Distance to Border of the DAPI-Intense Regions | µm | The mean euclidean distance of all the pixels of all the DIRs in the DIRs mask to the nucleus mask border |
| number of DIRs | Number of DAPI-intense Regions | - | The number of segmented DIRs using Watershed algorithm |
| largest slide perimeter | Largest Slide Nuclear Perimeter | µm | The perimeter of the nucleus mask in the largest 2D slide |

| | | | |
|---|---|---|---|
| largest slide area | Largest Slide Nuclear Area | $\mu m^2$ | The area of the nucleus mask in the largest 2D slide |
| largest slide major axis length | Length of Major Axis of the Largest Slide | μm | The length of the major axis of the ellipse that has the same normalized second central moments as the nucleus mask of the largest 2D slide |
| largest slide minor axis length | Length of Minor Axis of the Largest Slide | μm | The length of the minor axis of the ellipse that has the same normalized second central moments as the nucleus mask of the largest 2D slide |
| largest slide roundness | Largest Slide Nuclear Roundness | - | The roundness of the nucleus mask in the largest 2D slide relative to a circle. Computed as indicated in equation 2 below. |
| largest slide EOP | Largest Slide Nuclear Excess of Perimeter | μm | The excess of the perimeter of the nucleus mask in the largest 2D slide compared to an ellipse with the same major and minor axis lengths. Computed as indicated in equation 3 below. |
| intensity 0μm-0.5μm | Mean Intensity within 0μm-0.5μm along the Nuclear Border | - | The mean intensity value from all the pixel within the range of 0μm-0.5μm from the nuclear border in 3D |
| intensity 1μm-1.5μm | Mean Intensity within 1μm-1.5μm along the Nuclear Border | - | The mean intensity value from all the pixel within the range of 1μm-1.5μm from the nuclear border in 3D |
| intensity ratio | Nuclear Intensity to Border Ratio | - | The ratio of the mean intensity in the 1μm-1.5μm distance interval from the nuclear border to the mean intensity in the 0μm-0.5μm distance interval from the nuclear border |

**Equation 1.**

$$sphericity = \frac{\sqrt[3]{36\,\pi\,V^2}}{A_{surface}}$$

Where V denotes nuclear voxel volume in 3D and $A_{surface}$ is the nuclear surface area.

**Equation 2.**

$$roundness = 4\,\pi\,\frac{A}{P^2}$$

Where P denotes the nuclear mask perimeter and A is the 2D nuclear area.

**Equation 3.**

$$EOP = \frac{P_N - P_E}{P_E}$$

Where $P_N$ denotes the nuclear perimeter and $P_E$ is the perimeter of an ellipse with identical major and minor axis lengths as the measured nucleus:
